# The impact of micro-habitat fragmentation on microbial populations growth dynamics

**DOI:** 10.1101/2024.04.05.588087

**Authors:** Dina Mant, Tomer Orevi, Nadav Kashtan

## Abstract

Microbial communities inhabit almost every habitat on Earth and are essential to the function of diverse ecosystems. Most microbial habitats are not spatially continuous and well-mixed, but rather composed, at the microscale, of many isolated or semi-isolated local patches, resulting in partitioning of microbial populations into discrete local populations. The impact of this spatial fragmentation on population dynamics is not well-understood. Here, we study how fragmentations affect the growth dynamics of clonal microbial populations and how dynamics in individual patches dictate those of the whole metapopulation. To investigate this, we developed the µ-SPLASH, a novel ecology-on-a-chip platform, enabling the culture of microbes in microscopic landscapes comprised of thousands of microdroplets, spanning a wide range of sizes. Using the µ-SPLASH, we cultured the model bacteria *E. coli* and based on time-lapse microscopy, analyzed the population dynamics within thousands of individual droplets at single-cell resolution. Our results reveal that growth curves vary dramatically with droplet size. While growth rates generally increase with drop size, reproductive success and the time to approach carrying capacity, display non-monotonic patterns. Combining µ-SPLASH experiments with computational modeling, we show that these patterns result from both stochastic and deterministic processes, and demonstrate the roles of initial population density, patchiness, and patch size distribution in dictating the local and metapopulation dynamics. This study reveals basic principles that elucidate the effects of habitat fragmentation and population partitioning on microbial population dynamics. These insights are imperative for a deeper understanding of natural microbial communities and have significant implications for microbiome engineering.

## Introduction

Most microbial habitats are not spatially continuous and uniform but rather spatially fragmented into distinct, isolated or semi-isolated micro-patches, each hosting a small local microbial community. These patches are often heterogeneous in their characteristics, for example they can differ in their size, their physicochemical conditions, and available resources^1–4^. Typical examples of fragmented microbial habitats include microdroplets on leaf surfaces^5–11^, porous media in soil^12–16^, glands and hair follicles on the human skin^17,18^, crypts, digested food or mucus encapsulation in the gut^19–22^, and sinking particles in the ocean^23–25^.

To study the dynamics of microbial communities and populations, researchers often collect samples from macro-habitats such as a whole plant or leaf, a soil grain, a teaspoon of fecal matter, or a milliliter of ocean water, and then extract and analyze the microbial communities found within them. These studies typically average, or aggregate, the dynamics across communities or metapopulations^26,27^. Importantly, this approach either eliminates or ignores, the existing partition into local sub-populations, thereby neglecting the nuanced dynamics occurring within each micro-patch. Furthermore, the standard laboratory setup – relying on well-mixed, uniform liquid cultures in tubes or multi-well plates – stands in contrast to the patchy, heterogeneous nature of real-world microbial environments characterized by many localized, small populations^11,18,28–31^. These approaches overlook the intricate ways these local-populations contribute to the overall macro-population (metapopulation) dynamics. Consequently, there is a gap between how we measure, model and understand population dynamics and the actual manifestation of these dynamics within natural microbial communities.

The question how habitat fragmentation affect community and population dynamics has been more often studied in macro-ecology, including studies of landscape ecology^32,33^ that seek to understand how habitat structure and fragmentation affect biodiversity^34–38^ and population dynamics^39–41^ on both ecological and evolutionary timescales^39–44^. A substantial body of research shows that spatial heterogeneity affects species interactions, as well as the resulting population dynamics in both micro- and macro-ecology^44–53^. For example, partition of the metapopulation into local sub-populations has been found to affect competition and co-existence^15,54–57^, and to render inter-species interactions more stochastic, in particular in environments with sparse populations and low census numbers. This effect highlights the critical role of habitat fragmentation and population partitioning in shaping microbial population dynamics, and the resulting community composition and diversity^51^, as evidenced by recent studies, including those by Wu et. al^58^ and Batch et. al^59^.

Therefore, recognizing the intricate microscale structure of microbial habitats unveils a primary challenge in microbial ecology: bridging the gap between the micro and macro scales^60,61^, a key aspect that our current study seeks to address. Spatial variation in the microscale is a common feature of most microbial habitats. Consequently, microorganisms that can be just few tens of microns apart but reside in different ‘patch’ (e.g., microdroplet) can experience very different biotic and abiotic conditions. Thus, to accurately interpret population dynamics observed at the macroscale, it is crucial to develop experimental tools that capture the fragmented nature of the habitat under study. These tools should facilitate the monitoring of growth and dynamics of microbial sub-populations within their specific local micro-habitats. Alongside such experiments, the development of mechanistic and quantitative models and theories is necessary. Such advancements would enable the integration of insights from both micro and macro scales ^61–63^, aiding in the explanation of experimental observations. One promising approach is the development of ecology-on-a-chip platforms^30,64–67^. These platforms provide a replicable and controllable means to capture essential characteristics of natural habitats, thereby bridging the micro-macro divide in microbial ecology.

In this study, we adopted a systems microbial ecology approach combining experiments in a simplified *in vitro* system and computational modeling. Our goal was to systematically and quantitatively investigate the impact of micro-habitat fragmentation on population growth dynamics of the simplest case of clonal *E. coli* populations. Specifically, we aimed to understand how the dynamics of local sub-populations in many individual droplets, that vary in size, collectively modulate the metapopulation dynamics across the entire macro-habitat. We hypothesize that growth curves, along with their underlying kinetic variables, can be very different among patches that vary in their size.

To accomplish this, we developed a novel ecology-on-a-chip experimental platform, the µ-SPLASH chip, (Fig. 1). The system incorporates an airbrush that generates a jet of micro-sized droplets originating from preloaded suspensions of fluorescently-tagged bacteria. The sample is sprayed onto a glass-bottom surface, creating a microscopic waterscape of thousands sessile microdroplets ranging from picoliters (pL) to tens of nanoliters (nL) on a single ∼1 cm^2^ chip. Using microscopy and time-lapse imaging we were able to track *E. coli* growth curves in thousands of individual microdroplets at single-cell resolution.

**Figure 1.**
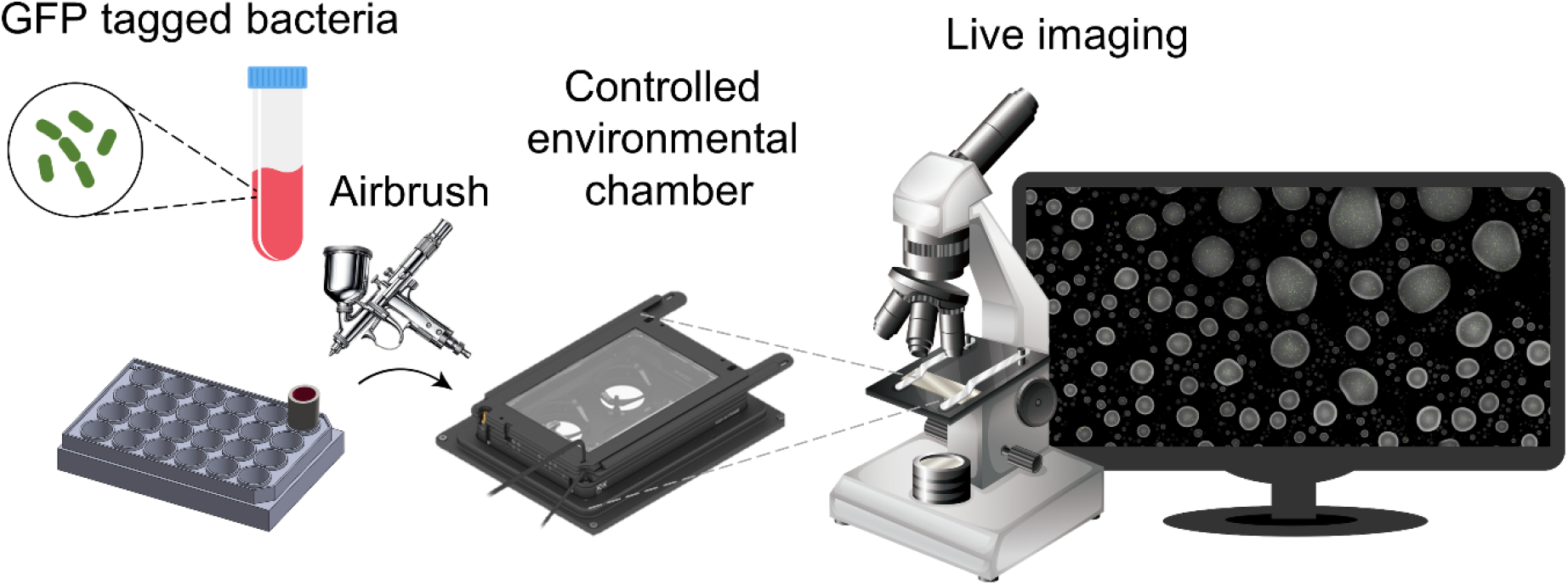
The µ-SPLASH experimental platform. Fluorescently tagged bacteria are sprayed onto a multi-well plate using a commercial airbrush. The plate is placed in a stage-top controlled environmental chamber set at 28°C and approximately 100% RH. Hourly imaging is conducted using a Nikon Eclipse Ti-E inverted microscope.

Our results reveal that, even within clonal bacterial populations, growth dynamics can significantly differ across patches of varying sizes. This is due to both stochastic and deterministic processes. Growth rate, reproductive success and the time to reach stationary phase show complex patterns that correlate with patch size. Therefore, integrating the growth dynamics of local populations from individual patches into the broader metapopulation dynamics, underscores how patchiness and the distribution of patch sizes, can profoundly affect the overall growth dynamics and yield of the metapopulation within a macro-habitat.

## Results

### The µ-SPLASH: an ecology-on-a-chip platform for studying population dynamics in fragmented microbial habitats

To study the impact of habitat fragmentation on population dynamics, we developed the µ-SPLASH system (Fig. 1). This system relies on an airbrush spraying of a solution containing bacterial cells onto a glass-bottom multi-well plate. This technique generates thousands of microdroplets with varying volumes ranging from 1 pL to 100 nL, all within a single chip (we define a ‘chip’ as a section of a single well that was scanned under the microscope).

After spraying, the well-plate is sealed and placed under the microscope within an on-stage controlled environmental chamber. This setup ensures that each droplet maintains its size throughout the duration of the experiment. Using a fluorescence microscope, each chip underwent hourly imaging of large surface area of approximately 6 mm x 6 mm for a duration of 24 hours. These scans produced high-resolution time-lapse imaging of thousands of microdroplets per chip, enabling the tracking of population dynamics at single-cell resolution (Fig. 2). The extensive gained dataset provided a robust foundation for assessing bacterial populations dynamics within thousands of individual droplets at single-cell resolution.

**Figure 2.**
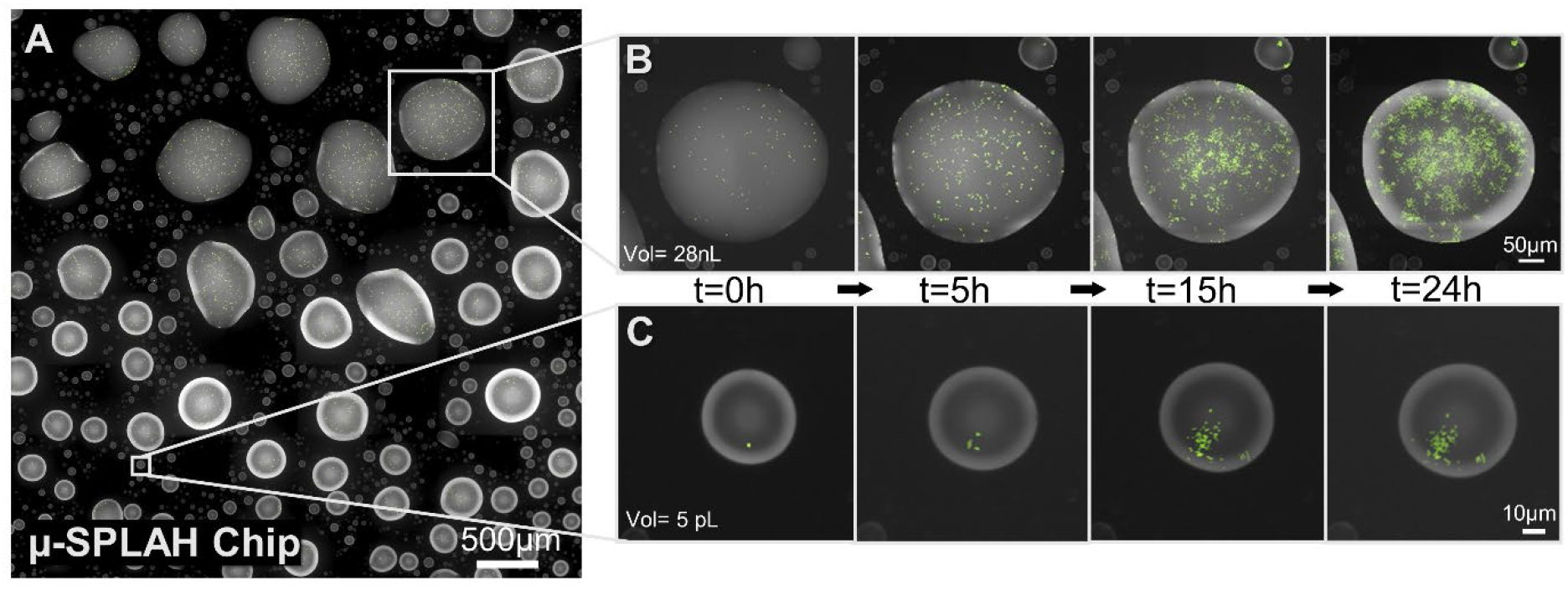
*E. coli* growth on the µ-SPLASH chip. (A) A representative section of a µ-SPLASH chip showing microdroplets spanning a wide size range (from 1pL to 100 nL). **(B-C)** Time-lapse series depicting *E. coli* growth (in green) within a large (B) and a small (C) droplet at t=0, t=5, t=15 and t=24 h. Cell population in droplets ranged between few cells in small droplets to thousands of cells in larger ones.

In the current study we analyzed 11 µ-SPLASH chips inoculated with *E. coli* cells growing in minimal media. For clarity and ease of presentation we first describe the results of a single representative chip (C6HD). Subsequently, we present the aggregated results of all 11 chips and discuss the overall findings.

### Seeding of bacterial cells into droplets is stochastic

We initially aimed to assess the distribution of droplet size on the chip. To accomplish this, we scanned an area of approximately 6mm x 6mm of the µ-SPLASH chip and utilized the Alexa (a soluble fluorescent dye that was used to mark the droplets) channel to segment the droplets. Next, we converted the droplet area, determined from the segmentation, into volume, based on the contact angle (see Materials and Methods). Our analysis revealed that droplets spanned a wide volume range, from 10^3^ µm^3^ (1 pL) to 10^8^ µm^3^ (100 nL; Fig. 3A). Not all droplets contained bacteria. Out of the total number of droplets on the chip, a subset contained at least one bacterial cell at time t=0 h (404 droplets out of a total of 1111 droplets on the chip). As expected, we observed that the larger the volume the higher was the likelihood that it contained cells at t=0 h. Specifically, most small droplets (10^3^-10^5^ µm^3^) did not contain any cells, some of them contain just one single cell (e.g., Fig. 2C). Virtually all droplets larger than 10^5^ µm^3^ harbored bacteria with up to thousands of cells in the largest droplets (Fig. 2B and 3A).

**Figure 3.**
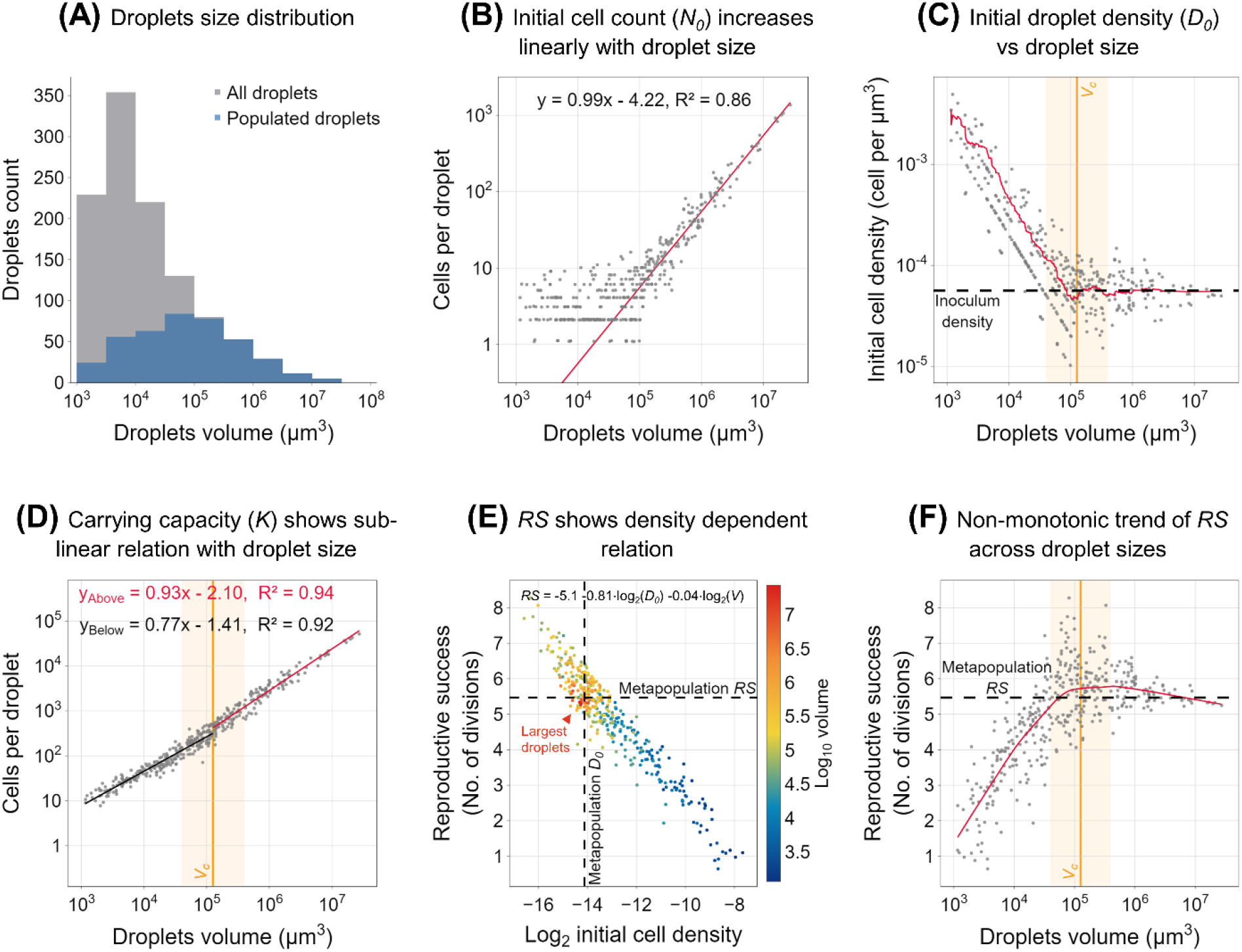
Initial cell numbers, carrying capacity and reproductive success show distinct patterns as a function of droplet volume. (A) The distribution of droplet volumes on a single chip. Gray represent all droplets, while blue indicates droplets populated with bacteria. **(B)** Bacterial cell count at t=0 h (*N_0_*) in individual droplets as a function of droplet volume (log-log plot). *N_0_* shows a linear relationship with volume. **(C)** Initial cell density at t=0 h (*D_0_*). The red line denotes a moving average of log-transformed values (window=50). The black dashed line represents the metapopulation cell density, i.e., the overall initial cell counts within all droplets divided by the total volume of all droplets. The vertical orange color represents the critical droplet volume (*Vc*), above which the mean cell density converged to the metapopulation cell density. The background orange color represents the intermediate droplets range (10^4.5^-10^5.5^ µm^3^). **(D)** The relation between carrying capacity (*K*) and droplet volume (log-log plot), *K* shows two sub-linera phases with different exponents. **(E)** Reproductive success (*RS*; the number of cell divisions per founder cell) as a function of droplet density and volume. Color map represents droplet’s volume. Note a weak yet significant negative contribution of *V* to *RS*, as observed in a vertical downward gradient for equal-density droplets (see e.g., around log_2_*D_0_*=-15), as well a small negative coefficient (*B*=-0.04) in the multiple regression model: *RS* = -5.1– 0.81⋅log_2_(*D_0_*) – 0.04⋅log_2_ (*V*). **(F)** Reproductive success (*RS*) as a function of droplet volume. The red line represents the locally weighted scatterplot smoothing (LOWESS, frac=0.4) trendline. The background orange color represents the intermediate droplets range around *Vc* (10^4.5^-10^5.5^ µm^3^), where *RS* was maximal. The dashed black line represents the metapopulation *RS*.

This observed pattern aligns with the expectation that cells were stochastically distributed between droplets in a proportion to droplet volume, thus following a Poisson process. Indeed, the initial number of cells per droplet (*N_0_*) showed a strong linear relation to droplet volume (*V*), reflected in an exponent ≈1 in a log-log plot (Fig. 3B, regression coefficient of 0.99 ± 0.03 for droplets with *V* > 10^5^ µm^3^, R²=0.86). The coefficient of variation (CV) of *N_0_* in the small droplets (10^3^-10^5^ µm^3^) was high (0.52), reflecting the stochastic nature of bacterial seeding in these smaller volumes. In larger droplets (10^5^-10^8^ µm^3^), however, there was a shift towards a more deterministic pattern with lower CV (0.36; Fig. 3B).

Next, we focused on analyzing cell density, a critical variable influencing population dynamics, in each individual droplet. We calculated the initial cell density (*D_0_*) at time t=0 h in all droplets that contained at least one cell at that time point. Interestingly, we observed that cell density in the smallest populated droplets was up to eighty times higher than the mean overall density of the entire metapopulation. The expected cell number in the smallest droplets, based on their volume multiplied by the cell density in the spray, was much lower than 1. Consequently, most small droplets contained zero cells; however, the small fraction of droplets that incidentally seeded with one, or few cells, exhibited very high density. As anticipated, cell density exhibited an exponential decrease with droplet volume, until reaching a density comparable to that of the sprayed suspension (Fig. 3C). Specifically, the mean *D_0_* decreased with volume in droplets of <10^5^ µm^3^ and then virtually converged towards the metapopulation density (*D_0_* = 5.63e-5 µm^-3^) at a volume that we term *Vc* (critical volume; here *Vc*=10^5.1^ µm^3^). These findings underscored the considerable heterogeneity in initial cell density values, where both the mean and the variance of *D_0_* varied with drop size.

### The carrying capacity shows two sub-linear phases relative to droplet volume

The carrying capacity is the maximal number of cells a given droplet can support. In our experiments, it was equivalent to yield or productivity, as growth curves reached saturation within the experimental timeframe. We opted to use the term carrying capacity (*K*), commonly employed in population dynamics models. Practically, we calculated *K* as the average cell count in the final four hours of our 24-hour experiment. This time period was chosen as most growth curves had plateaued by approximately 20 hours, indicating that the carrying capacity had likely been attained. The carrying capacity exhibited, at a broader view, a power-law relation with droplet volume (Fig. 3D). However, upon detailed examination it becomes evident that the behavior of K encompasses two phases: for droplets smaller than *Vc*, the slope is lower (0.77 ± 0.01), while for droplets larger than *Vc*, the slope is higher, yet, remains slightly smaller than 1 (0.93 ± 0.02). A slope lower than 1, observed in both of these phases, suggests a sub-linear, and not perfectly linear, relationship between *K* and *V*. If nutrient availability were the sole limitation, a perfectly linear relationship would have been expected for the large droplets (where *V* > *Vc*; the initial cell density is equal to the inoculum cell density). This indicates that nutrient availability significantly influences *K* (and probably is the most important factor), yet other unidentified factors further constrained growth.

### Reproductive success depends mostly on initial cell density but also on volume

Reproductive success (*RS)* is defined as the number of cell divisions per founder cell by time *t*. Over the entire experiment duration, the reproductive success of each local population can be calculated as *RS* = log_2_ (*K* / *N_0_*), where *K* is the carrying capacity and *N_0_* is the initial number of cells per droplet. Using a multiple linear regression model for reproductive success in individual droplets, *RS* = *A* + *B*log2(*D_0_*) + *C*log_2_(*V*), we found a very good fit (R^2^ = 0.91, see Methods, Supp. Table S1) indicating that *RS* is primarily influenced by the initial cell density (*B* = -0.81) and to a lesser extent by droplet volume (*C*=-0.04) (Fig. 3E). Reflecting on the dual-phase pattern of *K* observed in Fig. 3D, this model, together with the initial cell density pattern (Fig. 3C), can explain the two phases. For *V* < *Vc* the decrease in *D_0_* predominates, resulting in a lower exponent of ≈0.77. For *V* > *Vc*, on the other hand, where *D_0_* already converged at the inoculum density, droplet volume exerts only a weak negative effect, leading to a higher exponent, yet somewhat smaller than 1 (≈0.93).

### Reproductive success showed a non-monotonic function of droplet volume

Interestingly, we observed that the behavior of droplet *RS* as a function of droplet volume exhibited a non-monotonic pattern (Fig. 3F). Specifically, *RS* peaked at intermediate-sized droplets around the critical volume *Vc* (10^4.5^–10^5.5^ μm^3^), rather than in the largest droplets. Several factors account for this non-monotonic pattern. The first and most important one is the dependency of *K* on the initial cell density. Small droplets (*V* < *Vc*) had low *RS* that increases with volume. Despite many small droplets being empty, those with bacteria had initial cell densities much higher than the average initial cell density (Fig. 3C). Thus, even a single bacterium in such a small droplet falls near the carrying capacity, leading to low *RS* values. In contrast, in large droplets (*V* > *Vc*) the initial cell density – approximately equal to the inoculum cell density – falls below the carrying capacity, thus allowing population growth, as reflected by high *RS* values. Droplets in the intermediate volume range around *Vc* (10^4.5^-10^5.5^ μm^3^) showed pronounced variance in initial cell density. Due to the stochastic nature of droplet seeding with cells, some droplets in this size range exhibited very low initial cell densities, while others exhibited relatively high initial cell density values (Fig. 3C). This high stochasticity in initial cell density in the intermediate-sized droplets led to many droplets with very high *RS*, much higher than that of the largest droplets. This results in an overall increase in the fraction of the population residing in intermediate size droplets and a decrease in the fraction that resides in small and large droplets (Supp. Fig. S1).

Another important feature of the observed *RS* pattern is its slight but significant overall decline from intermediate-sized drops to large droplets (Fig. 3F). This decline can be explained by the weak negative effect of volume on *RS* (Fig. 3E) reflected by observation that *K* increases sub-linearly (and not perfectly linearly) with volume (exponent=0.93, for *V* > *Vc*) (Fig. 3D). Indeed, the intermediate-sized droplets had an average *RS* of 5.69 ± 0.89 (mean ±SD) which was significantly higher (one-sample t-test, *P* = 0.002) than the overall metapopulation *RS* (5.46), calculated as *RS* = log_2_(sum (*K_i_*) /sum (*N_0_*)) (dashed line Fig. 3F). These observations underscore the heterogeneity of *RS* across droplets, its non-monotonic dependence on the volume, and how *RS* in individual droplets can deviate from the entire metapopulation *RS,* that is typically measured in lab experiments and field studies.

### Growth curves in individual droplets vary with droplet volume

To gain further insights into the growth patterns within each droplet, with respect to droplet size, we categorized the populated droplets into five logarithmic volume bins: 10^3^-10^4^, 10^4^-10^5^, 10^5^-10^6^, 10^6^-10^7^, and 10^7^-10^8^ µm³. In this chip, the total number of 404 populated droplets distributed into these bins as follows: 80, 148, 131, 40, and 5, respectively. For each bin, we plotted the mean ± SD growth curve over all droplets of that bin (Fig. 4A). These curves provide a general view of how growth dynamics vary across different droplet sizes. The growth curves differed from the average growth curve (averaging across all drops; Fig. 4A dashed line) or the aggregated growth curve of the entire metapopulation (Fig. 4A solid line). To visually compare the shapes of the growth curves across volumes, we normalized the growth curves of each droplet to its respective carrying capacity (*K*) (Fig. 4B). Several trends can be visually identified by examining this plot. First, the smaller droplets (10^3^ – 10^4.5^) started at higher ‘relative densities’ (i.e., *N_0_*/*K*) and approached *K* (carrying capacity) relatively early. In other words, the initial density in smaller drops lies closer to the density at carrying capacity (*K*). Second, the intermediate size droplets around the *Vc* (10^4.5^ – 10^5.5^) presented high variability in their growth curve shapes, and also approached *K* at various times (green-yellow). Third, the largest droplets (red) showed fairly uniform behavior and represented a mediocre growth curve. In summary, growth curves varied significantly across the metapopulation yet, cluster around each specific droplet size range.

**Figure 4.**
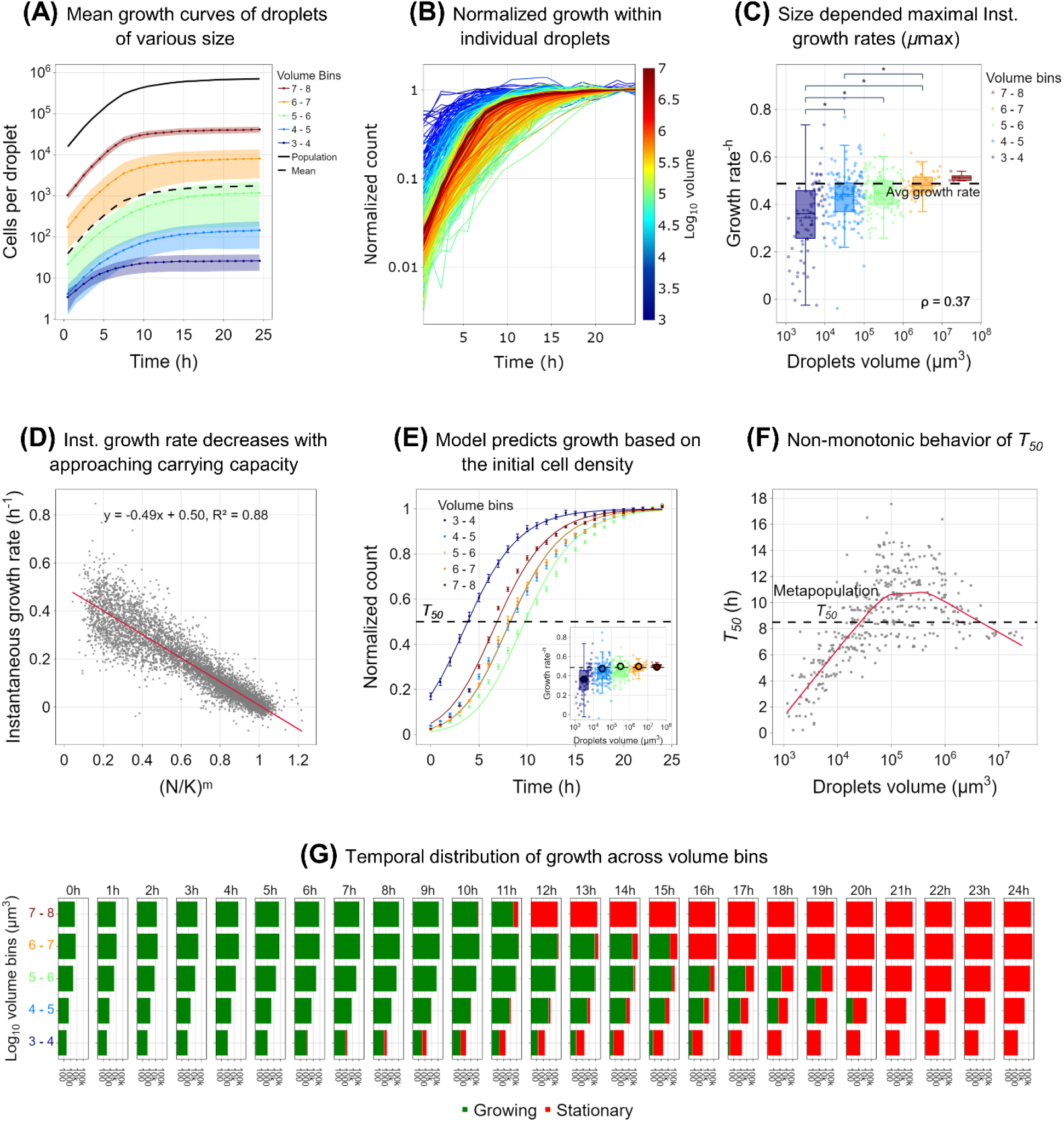
Growth curves, growth rates and time to approach the carrying capacity vary with droplet size. (A) Growth curves of the *E. coli* populations on the µ-SPLASH, represented by the mean cell count per droplet volume bins (log_10_ volume) over time (0-24 h). The shaded color of each curve represents the standard deviation. The unshaded dashes lines represent the overall metapopulation (upper) and the mean of all the droplets (lower). **(B)** Normalized growth curves within all populated individual droplets (404 droplets). Each droplet is colored based on the droplet’s volume. Y-axis in log scale. **(C)** Maximal instantaneous growth rate (*μ*_max_) of each droplet. The boxplot represents the *μ*_max_ distribution of the droplets within each volume bin, colored based on the volume bins. The dashed line represents the overall metapopulation *μ*_max_. A permutation tests (n=1000) was used to test statistical significance between the bins’ means. ρ represents Spearman rank correlation coefficients (P-value<3.2e-14). **(D)** Instantaneous growth rates against the ratio between bacterial cell numbers and carrying capacity in the power of ‘*m*’, the deceleration parameter. Linear relationship represented by OLS regression line. **(E)** Normalized growth curves for each volume bin (symbols and error bars represent Mean ± SE). The line represents the generalized logistic growth model for each volume bin, based on the median *r* and *m* values of all droplets. **(F)** Time to reach 50% of the carrying capacity (*T_50_*) as a function of droplet volume. The red line represents the locally weighted scatterplot smoothing (LOWESS) trendline. The dashed line represents the metapopulation *T_50_*. **(G)** Bar chart representing time-lapse of the overall population dynamics and state (growing in GREEN; stationary in RED) over time in each bin. The bar width represents the overall cell numbers (in log scale) in each drop-volume bin over time.

### Instantaneous growth rates vary with droplet size

To better understand the aforementioned pattern and the effect of drop size on bacterial growth, we analyzed the kinetic variables dictating these growth curves. First, we calculated the instantaneous growth rates (*μ*) in each droplet over the course of the experiment. To derive μ, we performed linear regressions on the natural log number of cells within a moving time window of four hours. The slope from each regression period represented μ for that time. Generally, *μ*_max_, defined as the highest μ, was observed at the first 4-hour window and decreased with time, until the population within each droplet approached the carrying capacity and growth rate declined to zero (Supp. Fig. S2). It is worth noting that in our experiments we did not observe a ‘lag phase’, which is often seen in bacterial cultures. This absence is likely due to the fact that cells were growth under similar conditions in the pre-cultures and were transferred from mid-log phase. In addition, there was a time gap of about 1 hour between spraying the cells on the chip and the first imaging time point. Interestingly, we observed that *μ*_max_ increased with droplet size (Spearman correlation r=0.37) (Fig. 4C). Specifically, *μ*_max_ in the smallest volume bin (10^3^-10^4^) (*μ*_max_ =0.34 ± 0.15 (h^-1^) mean ± SD), was significantly lower than *μ*_max_ in the 10^4^-10^5^, 10^5^-10^6^, 10^6^-10^7^ bins (0.43 ± 0.11, 0.45 ± 0.07, and 0.49 ± 0.05, respectively) (Fig. 4C). We note that small sized droplets presented a higher variance in *μ*_max_ which could stem from phenotypic variation^59^ or higher noise due to small cell numbers^59,68^.

### Density-dependent growth rate analysis based on the generalized logistic growth model

Next, we fitted the observed pattern of growth rates as a function of droplet size, to a well-established model for population dynamics, the generalized version of the logistic growth model^69^. This model takes into account the constraints on population increase due to density dependence and has been demonstrated to effectively describe bacterial growth^69–71^. This generalized logistic growth model serves as a fundamental tool in comprehending population dynamics, particularly in understanding how growth rate is influenced by density. The equation of this model is expressed as:

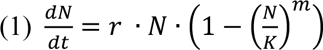

where *t* is time, *N* = *N*(*t*) is the cell number at time *t*, *r* is the intrinsic growth rate, *K* is the carrying capacity and *m* is a deceleration parameter. This model is a similar to the classic logistic growth model^72^ but it includes an additional parameter known as the deceleration parameter (*m*).

Applying this model to our data, we fitted the experimental growth curves within each individual droplet and extracted the median *r* and *m* of all droplets on the chip (*r*=0.55, *m*=0.61) (Supp. Fig. S3). Using the fitted median *m*, we next examined the density dependence of the instantaneous growth rates *μ(t)* with 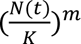. A strong linear decline (R =0.88) was observed, indicating a clear density-dependent growth relationship pattern (Fig. 4D). This trend suggests that growth rate declines as cell density in the droplet increases, which plays a crucial role in modulating growth rates within individual droplets.

Next, we aimed to explore whether the differences in the initial cell density across droplet size could account for the variations in *μ*_max_. To investigate this, we fitted the model to the mean normalized growth curves of droplets grouped by their size, using a single set of parameters: the median fitted *r* and *m* values. The predicted growth curves by the model, broadly agree with the experimental data (Fig. 4E). This comparison highlighted that the differences in growth curves across volume bins could primarily be attributed to initial cell density. Since the initial density in small droplets was higher than the metapopulation *D_0_* (Fig. 3C), their *μ*_max_ was relatively low. In contrast, larger droplets with *D_0_* closer to the metapopulation *D*, demonstrated higher *μ*_max_ values (Fig. 4C). Indeed, when we use the model to predict the observed *μ*_max_ by plugging the median *r* and *m* values into the model, along with the mean initial ‘relative density’ (*N_0_*/*K*) of each size bin, we observed a good agreement with the measured *μ*_max_ values and the overall trend (Fig. 4E inset). These results reinforced the notion that initial cell density in individual droplets plays a crucial role in determining growth rates in individual droplets.

### Characteristic growth time (*T_50_*) varies among droplets and exhibits a non-monotonic pattern

Another key kinetic variable of growth curves is termed *T_50_* – the time to reach 50% of the carrying capacity. *T_50_* is a function of the initial density, growth rate change over time, and the carrying capacity. Our analysis of the average growth curves, as illustrated in Fig. 4E, indicated that *T_50_* varied with droplet size. Importantly, we observed a non-monotonic trend in *T_50_* as a function of droplet volume. The intermediate-size droplets (10^4.5^-10^5.5^), exhibited the highest *T_50_* (Fig. 4F). This result is consistent with the observed pattern for reproductive success, wherein the intermediate-size droplets also demonstrated the highest reproductive success.

The non-monotonic behavior of *T_50_* as a function of droplet size, led to another important observation. We asked, at each time point, what fraction of the population was still growing (i.e., in exponential phase). To this end, for each droplet we determined the time it took to reach 80% of the carrying capacity as a proxy of the time of reaching the carrying capacity. Our temporal analysis of droplet populations (Fig. 4G) showcased the dynamics of bacterial growth as droplets progressed towards their carrying capacity. We find the visualization in Fig. 4G helpful in observing what fraction of cells are growing and which have reached stationary phase, over time and across droplet size bins. This bar chart revealed an apparent pattern: smaller droplets (10^3^-10^4^ µm³) stopped growing first, at t=∼7,8 h, followed by the largest droplets (10^7^-10^8^ µm³) which stopped at t=∼11-12 h. The intermediate-sized droplets stopped growing over a wider time window at t=13-20 h. Interestingly this highlights another non-trivial observation, as it indicates that at various time points, the actively growing cells—or from another perspective, the cells that have entered the stationary phase—reside in different droplet-size bins. This has significant ecological implications, such as response to antibiotics. Moreover, the sustained growth within intermediate-sized droplets, along with their higher *RS*, suggests that their relative contribution to the overall metapopulation increases over time (Supp. Fig S1). These unique dynamics may significantly affect natural selection and population distribution and interactions within patchy heterogeneous environments.

### Understanding how growth dynamics in individual droplets dictates the dynamics of the overall metapopulation

The results presented in the previous sections demonstrate significant variations in population dynamics across droplets of different sizes. We showed that reproductive success, growth rates and *T_50_* can be very different in droplets of different sizes. In this specific chip, reproductive success in some of the smallest droplets showed values close to just one cell division, while some intermediate-size droplets had a *RS* of 8 cell divisions (Fig. 3E-F).

Although there are stochastic factors affecting *RS*, it showed a clear dependency with droplet size. Thus, the overall *RS* of the whole metapopulation depends on the distribution of droplet sizes. Moreover, the growth rates differences across droplet size range further emphasize the role of droplet size distribution (Fig. 4C) on building up the average metapopulation growth rate. Droplet size distribution also significantly influences *T_50_* and the proportion of the population in either growth or stationary phase, at any given time (Figs. 4F, G). Collectively, these findings highlight the profound effect of droplet size distribution on the overall metapopulation dynamics (Fig. 4A).

### Comparative analysis across 11 μ-SPLASH chips

Thus far, we reported the results obtained from a single representative chip. Next, we wanted to ask (i) how robust are the observed patterns, and (ii) since the initial cell density seems to be a central variable, we wanted to know how the initial cell density (of the inoculum sprayed solution) affects the results. We present here results of 11 chips: five of them were inoculated with an initial sprayed suspension of OD = 0.01 (Low density; LD) and six with OD = 0.03 (High density, HD). The chip described in the previous section was seeded with OD=0.03 (chip ID C6HD). The total number of droplets analyzed in these 11 chips was approximately 9,000 in the LD scenario and 8,000 in the HD scenario (Supp, Table. S2). Overall, the drop size distribution was broadly similar across all chips (Supp. Fig. S4, Fig. 5A-B, Supp. Table. S2). Notably, in line with our expectations, the number of populated drops in the HD chips were higher (with a mean of 29.18% ± 0.51% of droplets containing bacteria in the HD scenario compared to 16.75% ± 0.4% in the LD scenario), and the peak of droplet size containing bacteria shifted towards smaller values (10^4.6^ µm^3^ in HD compared with the 10^5.2^ µm^3^ in LD).

**Figure 5.**
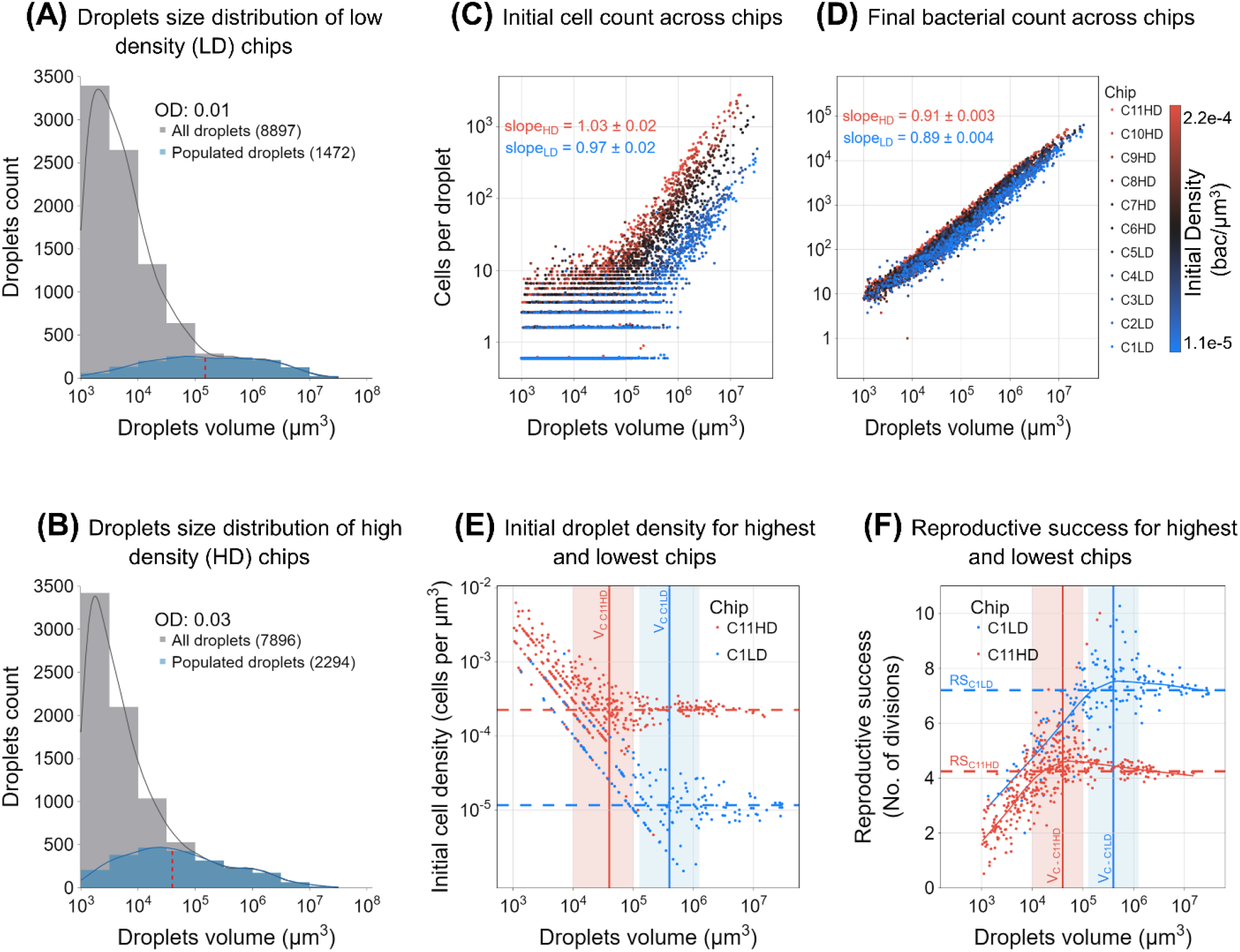
Joint-analysis of 11 µ-SPLASH experiments with increasing initial cell densities. (A, B) Aggregated droplet distribution in two initial sprayed ODs (0.01 and 0.03). In gray are all the droplets, and in blue are the bacteria-containing droplets only. Kernel density estimate (KDE) lines are shown. The red dashed lines indicate the median of the bacteria-containing droplets distributions. **(C)** Initial cell counts (*N_0_*) as a function of droplet volume for all 11 chips. Each color represents a different chip, while the color represents the initial cell density within each chip. Blue: the least dense chip; Red: the densest one. **(D)** Same as (C) for carrying capacity. **(E)** Initial cell density at t=0 h (*D_0_*) for the least and most dense chips. In blue, the least dense chip and in red the densest chip. The dashed lines represent the metapopulation initial cell density of each chip, i.e., the overall initial cell counts within all droplets divided by the total droplets volume. **(F)** Reproductive success (*RS*, mean number of divisions) as a function of droplet’s volume for the least and most dense chips. In blue, the least dense chip and in red the densest chip. Dashed lines represent the respective average reproductive success within each chip.

Although we used only two initial cell densities in the sprayed suspension, the sprayed cell density showed minor differences between chips. This is likely due to small differences in the preparation or other stochastic effects (such as minimal evaporation of droplets that occur between spray application and plate sealing) resulting in minor density differences, even within different wells sprayed with the same suspension. In fact, this feature proved to be advantageous as we captured even a wider initial density range which allowed a more fine-grain resolution (Fig. 5C). For the ease of visual presentation, we color-coded that data from these 11 chips based on their actual initial density (*D* = sum(*D_0_)* / sum(*V*)), assigning a color gradient with the lowest initial densities in blue and highest in red. The initial cell density in the highest density chip was ∼20 times higher than the lowest density one (Supp. Table. S2). The initial number of cells per droplet showed linear relationship with drop volume (exponent ∼1 in all chips, Fig. 5C, Supp. Table. S3), as expected. The differences in the initial cell densities were nicely reflected in the fitted line shifting upward with density. Moreover, we observed that with increasing initial cell density, there was a shift to the left in the stochastic range of initial cell numbers in the droplets.

Analysis of the individual droplet carrying capacities, in each of these chips, showed a consistent sub-linear relation, as was observed for chip C6HD (Fig. 3D), with an average overall exponent of 0.91±0.01 for HD chips and 0.94±0.01 for LD chips. *Vc* was also a function of initial cell density and decreased with density, thus affecting the biphasic K pattern (supp. Fig. S5, Supp. Table. S4). Interestingly, droplets with higher initial cell counts exhibited a significant increased carrying capacity for the same drop volume. Similarly to the results for chip C6HD, the initial density decreased with volume for the smaller droplets, while in the larger droplets the initial density converged to the mean initial density of the metapopulation. However, the mean overall density, affected the critical volume *Vc.* Thus, the transition point between the two trends mentioned above, shifted between different chips, representing different initial cell densities (Fig. 5E).

Next, we compared the behavior of reproductive success as function of droplet volume across chips (Supp. Fig. S6). The droplet *RS* derived from the chips with the lowest and highest initial densities (C1LD and C11HD, respectively) showed similar trends but differ significantly (Fig. 5F). As expected, the lower density chip showed a much higher average *RS* of 7.2 divisions compared to the chip with the highest density that showed 4.2 (Fig. 5F dashed lines). In the lowest density chip, the *RS* peaked in drop volumes between 10^5^ to 10^6^ µm³, while in the highest density chip, *RS* peaked in much smaller droplet volumes of 10^4^-10^5^ µm³ (Fig. 5F red and blue backgrounds). When comparing the aggregated data from all the LD chips with the HD ones, we observed a significant difference in the average RS (6.22 ± 1.47 in LD compared to 4.44 ± 1.27 in HD). Remarkably when we used a multiple linear regression model, we found a consistently similar coefficients across chips *RS* = -5.2 – 0.8log_2_(*D_0_*) – 0.01log_2_(*V*) (coefficient mean values; Supp. Table. S5).

The *T_50_* behavior across these chips also demonstrated variations with initial density, paralleling the trends observed in reproductive success. In chips with higher initial densities, the longest *T_50_* was exhibited in smaller droplets, starting at a volume ∼ 10^4.5^ µm³. Conversely, in the low-density chips, the highest *T_50_* was observed in larger droplets at a volume of ∼ 10^5.5^ µm³ (Supp. Fig. S7). While there were some variations in the growth curve shapes among the different chips, the underlying patterns observed remained robust (Supp. Fig. S8, S9). In some chips we noticed that cells in certain regions of the chip tended to organize as dense cell aggregates that dispersed at later times, which seemed to have an impact on the accuracy of the cell segmentation in these regions. This was reflected in less smooth growth curves in such droplets (Supp. Fig. S8, S9). Nevertheless, the overall patterns were consistent and robust. Growth rates were found to show similar density dependence across chips (Supp. Fig. S10). The non-monotonic patterns of *RS* and *T_50_* as a function of drop volume (Supp. Figs. S6 and S7), the increase in *μ*_max_ with volume (Supp. Fig. S11), the instantaneous growth rates (note some of the chips showed a late small peak which was correlated with cell dispersal at these times; Supp. Fig. S12), and the overall good fit of a generalized logistic growth (again, except the chips with cell aggregation and dispersal; Supp. Figs. S13), were repeatedly observed across all chips. Moreover, the proportion of the population in either growth or stationary phase at any given time presented a similar pattern across all the chips, as the intermediate-sized droplets took the longest time to reach the carrying capacity (Supp. Fig. S14). These consistent and robust patterns emerged across the 11 µ-SPLASH experiments reaffirms the validity and reliability of our findings.

### Computational model replicates the population growth dynamics in µ-SPLASH System

To validate our comprehension of the underlying principles governing cell growth dynamics in the µ-SPLASH system, we developed a simulation tool. The primary objective of the simulations was to accurately capture and replicate the growth patterns observed on the chip. The simulation was initialized with key parameters reflecting the empirical data, including the total volume of all droplets, the drop size distribution, and the initial cell density (see Methods). In these simulations, cells grew in individual droplets according to the generalized logistic model. Model parameters were those extracted from the experiments of the chip C6HD (Supp Table. S5).

In Figure 6 we present the simulation results based on measured parameters taken from C6HD, in comparison to the experimental results of this chip (inset). Our simulated droplet size distribution closely resembled the real one (Fig. 6A). The initial cell number showed linear relation with droplet volume (Fig. 6B), further indicating that droplet seeding with cells follow a Poisson process as in our modeling. Initial cell densities decreased with droplet size similarly to the experimental data and converged towards the inoculum density in the large droplets (>10^5^ µm^3^) (Fig. 6C). The carrying capacity reflected a similar function as in the real chip with two different phases shifting approximately at droplets of 10^5^ µm^3^, consistent with the sub-linear relationship observed in our experiments (Fig. 6D). The simulated reproductive success across varying droplet volumes demonstrated a non-monotonic relationship, in line with the empirical observations (Fig. 6E). The simulated normalized growth curves (large droplets in red,small droplets in blue) also showed a similar pattern to the empirical data: small droplets approached the carrying capacity the fastest, moderate droplets presented high variation in the time required to approach the carrying capacity, and large droplets approached the carrying capacity in intermediate timing (Fig. 6F). We noticed sized-dependent growth rate, that was similar to the empirical data, and showed higher growth rates in larger drops (Fig. 6G). Finally, the simulated *T_50_* also showed a non-monotonic pattern consistent with the experiments (Fig. 6H).

**Figure 6.**
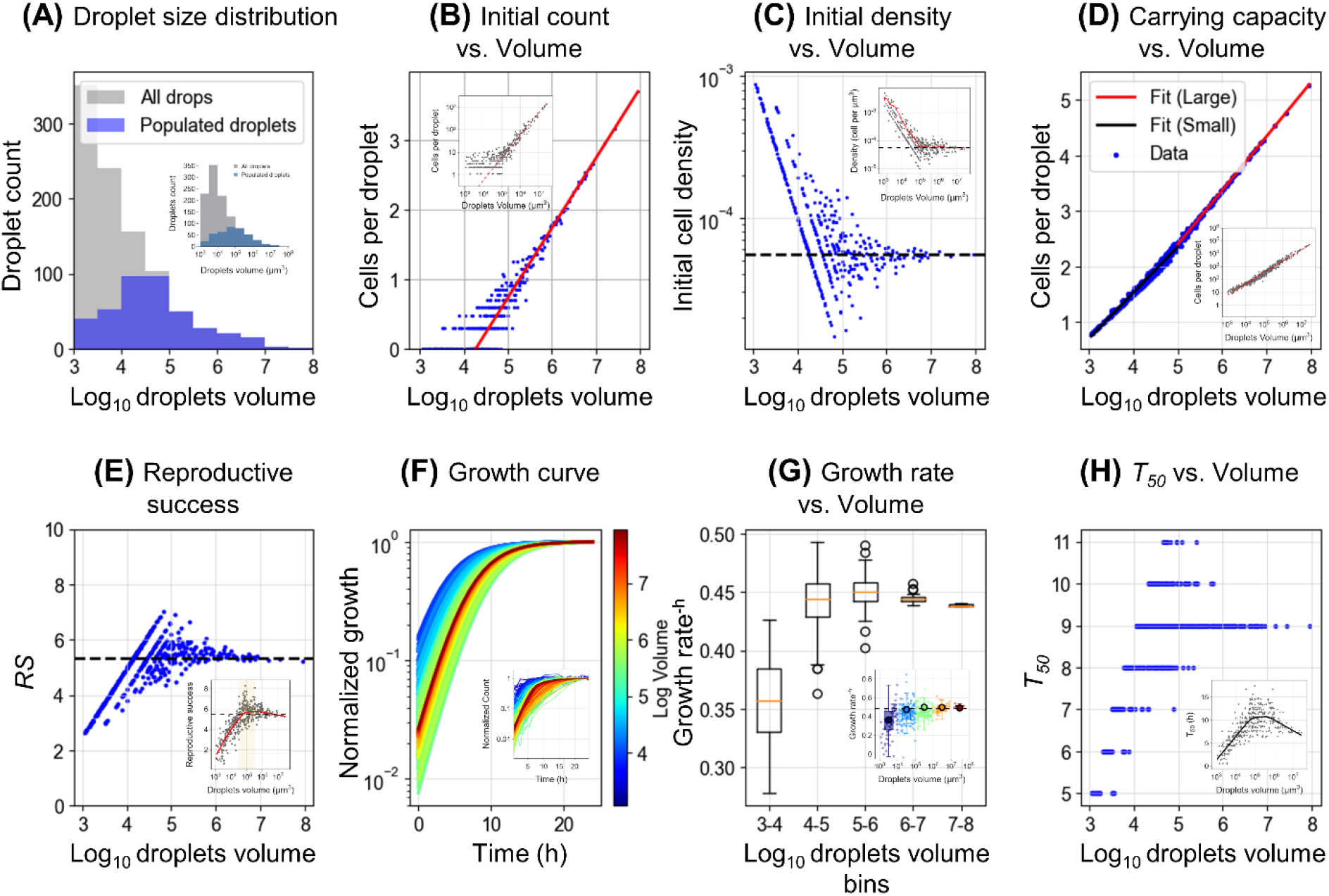
Simulated population growth dynamics of the µ-SPLASH. (A-E) Depict various aspects of the droplet generation and initial bacterial distribution, including histograms of droplet volumes, scatter plots of initial bacterial counts vs. droplet volume, the initial density across droplets, the relationship between droplet volume and carrying capacity and the reproductive success. Insets shows the experimental results **(F)** Shows normalized growth curves for bacteria within individual droplets over a simulated time period, color-coded based on the droplets log volume. **(G)** focus on the calculated bacterial growth rates across droplets categorized into log_10_ volume bins. Growth rates were determined by fitting a linear model to the log-transformed bacterial counts from t=0 h to t=4 h. **(H)** shows the time to reach 50% of the carrying capacity across different droplets volume.

### Exploring the impact of patchiness and drop-size heterogeneity on metapopulation yield and reproductive success

As the final chapter of this study, we aimed to extend our understanding of how patchiness and heterogeneity in patch (droplet) size influence key metapopulation dynamics metrics. First, we sought to investigate how the level of patchiness affects the overall metapopulation dynamics. We conducted simulations across a range of patch configurations, maintaining a constant overall volume but varying the number of patches from a single large droplet to up to 10^4^ small, equally-sized droplets (see Fig. 7A). For each patch configuration, we explored different initial cell densities, ranging from 10^2^ to 10^5^ cells per simulated chip.

**Figure 7.**
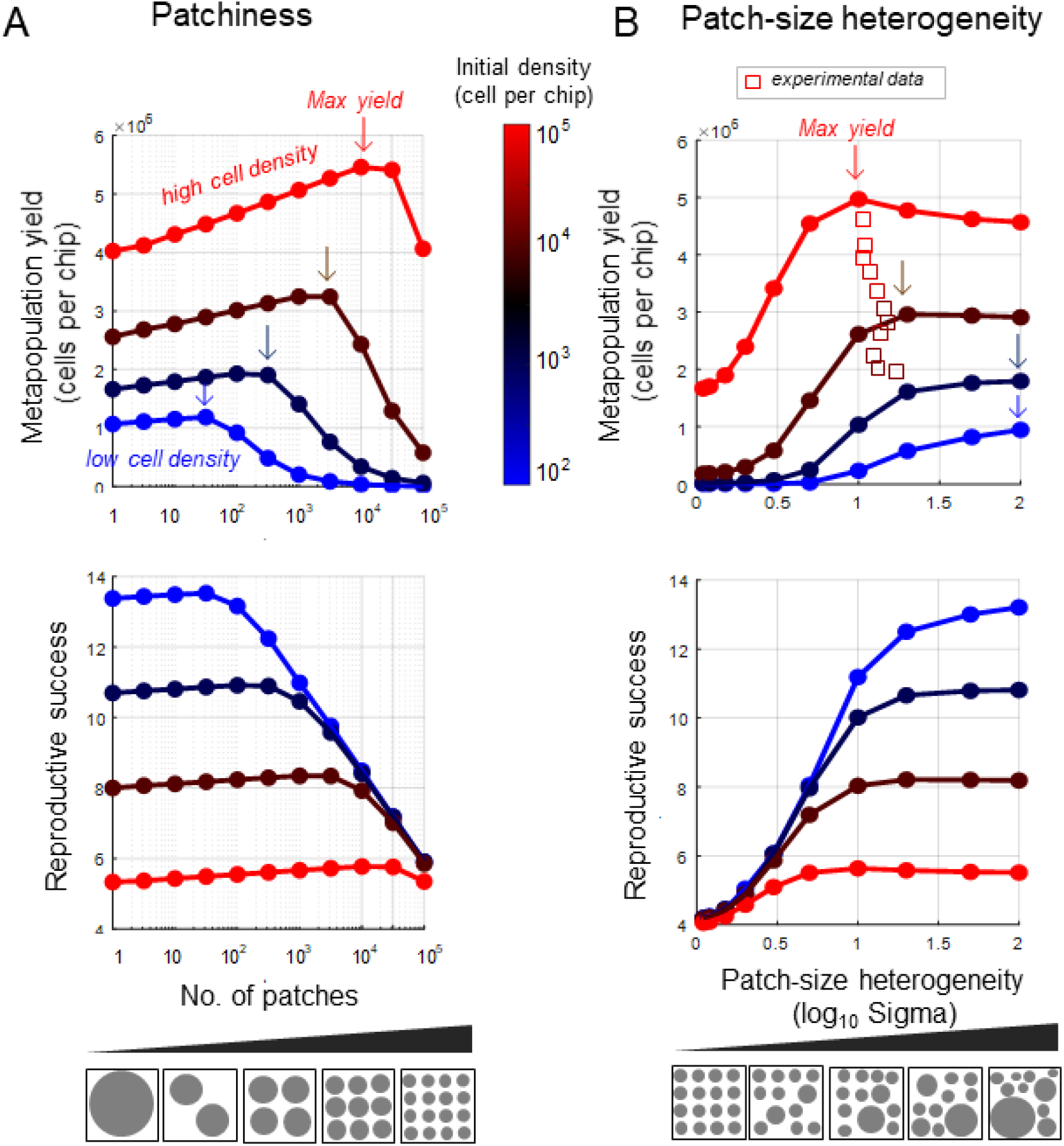
The impact of patchiness and patch size heterogeneity on simulated metapopulation yield and reproductive success. In red, brown, blue, and light blue are various initial cell densities (100,1000,10000,100000 per chip, respectively). The arrows denote the respective level in which each initial density level reached maximal yield. **(A)** The final metapopulation size (top) and the reproductive success (bottom) as a function of patchiness. Patchiness range varies from a single large patch to 100,000 small patches, summing up to the same total volume. **(B)** The final metapopulation size (top) and the reproductive success (bottom) as a function of heterogeneity in patch size. Patch size distribution was log normal with similar mean values (10^3.33^ µ^3^) and SD ranging from 1.1 to 100. Square symbols represent the experimental yield of the 11 chips normalized to equivalent volume of the simulated chip (10^9^ µ^3^) and their corresponding SD of their droplet size distribution; colors represent the initial cell density. Patchiness degree and heterogeneity is conceptually illustrated as circles with differing size and distributions.

Our simulations showed that the metapopulation yield (size of the metapopulation at t = 24 h) significantly vary with patchiness. The maximal yield was consistently observed at intermediate number of patches (Fig. 7A). Reproductive success (*RS*) followed a similar trend, with maximal number of divisions occurring at intermediate patch numbers. Notably, the level of patchiness for achieving the maximal yield and *RS* increased with initial cell densities (Fig. 7A). For the lowest initial cell density (10^2^ cells per chip) a maximal final population of ≈10^6^ cells was reached with 10 patches, corresponding to an RS of around 13 divisions. At moderate initial cell density (10^3^ cells per chip) a maximal yield was ≈2x10^6^ cells with approximately 300 patches, indicating an *RS* of about 7.5. In scenarios with higher initial cell densities of 10^3^ and 10^4^ cells per chip, the maximal yield was 3.2x10^6^ and 5x10^6^ cells, at 3x10^3^ and 3x10^4^ patches, respectively (Fig. 7A). These results show that yield depends on patchiness, and for a given initial cell density, the maximal yield is reached at different patchiness levels.

Next, we wanted to explore how patch-size heterogeneity affects the metapopulation yield and *RS*. To do so, we fixed a constant total volume while varying the variance (Sigma) of the droplet size distribution (Fig. 7B). As we kept the overall volume similar, the variance also affects the number of patches, but most importantly, altered the heterogeneity of droplet size. Our findings indicate that, for given initial cell density, the maximal yield and *RS* occur at different levels of heterogeneity. Yield differences in some cases exceeded an order of magnitude across waterscapes of varying heterogeneity. Generally, higher heterogeneity enhanced yield in lower initial densities, whereas more uniform waterscapes with many equally-sized droplets resulted in higher yield at higher initial cell densities.

These simulations enable us to significantly extend the range of parameters beyond what was feasible experimentally. They emphasize the profound effect of habitat fragmentation on population dynamics. Specifically, they highlight how varying levels of habitat patchiness can influence the metapopulation growth dynamics, demonstrating that an optimal degree of habitat fragmentation can maximize both population yield and reproductive success.

## Discussion

In this work, we studied the effect of a key characteristic of microbial habitats – its fragmentation into micro-patches of various sizes – on the growth dynamics of clonal bacterial populations. Using a newly developed ecology-on-a-chip platform, the µ-SPLASH, we cultured the model bacterium *E. coli* in artificial microscopic landscapes consisting of thousands of microdroplets, spanning a broad size range from picoliters to nanoliters. Our findings revealed significant variations in growth curves across droplets (patches) of different sizes. Specifically, we observed that growth rates varied with patch size and that reproductive success, and the time to approach the carrying capacity, peaked in intermediate-size patches. By combining µ-SPLASH experiments with computational modeling, we demonstrated that these observed patterns emerge from both stochastic and deterministic factors. Furthermore, our study illustrates how these factors modulate growth and population dynamics. Finally, we elucidated how growth dynamics in individual patches collectively influence the dynamics and yield of the entire metapopulation, underscoring the significance of patchiness and patch size distribution.

While growth rates generally increased with droplet size, reproductive success showed a non-monotonic pattern (Figs. 3,4), indicating that neither the smallest nor the largest droplets consistently yield the highest growth potential. Instead, droplets of an intermediate size—around a critical size determined by the inoculum cell density—often provide an advantageous environment that leads to higher reproductive success. This non-monotonic pattern results from two opposing trends: the reduced reproductive success in small droplets, which is attributed to higher initial cell densities that approach the carrying capacity, and the influence of a sub-linear relationship between droplet size and carrying capacity that becomes dominant in large droplets. As anticipated, the time to reach the carrying capacity mirrors the pattern of reproductive success, displaying a non-monotonic pattern with a peak at intermediate-size droplets.

We suggest that stochastic factors – key among them is the initial cell density in individual droplets – significantly affect the growth dynamics within each droplet. The variability in initial cell densities in individual droplets arises from the random allocation of bacterial cells into droplets, following a Poisson process. This randomness results in more variable cell numbers in small droplets which become more predictable in larger droplets. Consequently, smaller droplets have higher and more variable densities, while larger droplets have densities closer to the inoculum density, as shown in Figure 3C. This pattern is due to a critical droplet size threshold; below it, a substantial number of droplets remain unpopulated (Fig. 3A). Only a small fraction of droplets below this threshold size are stochastically seeded with one or a few founder cells (Fig. 3B), resulting in densities significantly above the expected level. Therefore, the initial cell density in these droplets is much higher than the inoculum density, greatly affecting growth rates and reproductive success. The pronounced negative density dependence of growth rates observed (Fig. 4D), along with the pattern of initial cell densities (Fig. 3C), explains the general increase in growth rates with droplet size (Fig. 4). The high cell density in smaller droplets give rise to lower reproductive success, as the initial cell count is already close to the carrying capacity (Fig. 3E). This inherent stochasticity of cell seeding into patches, often with only a few founder cells in smaller patches, is likely a common phenomenon across a wide range of natural microbial habitats – from the human gut to soil and plant surface micro-environments. Together with patch size heterogeneity is expected to have a profound impact on growth and reproductive success^73^. Thus, in essence, the stochastic nature of cell seeding is expected to play a pivotal role in the assembly and dynamics of microbial communities within fragmented habitats. It is also important to recognize that other stochastic factors, such as phenotypic variation, may also play a role, especially in smaller droplets with a few cells, particularly affecting interspecies interactions^59,68^.

Our findings further highlight the critical role that the source’s (e.g., inoculum) initial cell density plays in shaping population dynamics and yield. This conclusion is supported by our experimental data from multiple chips with varying initial cell densities (Fig. 5), as well as by our computational simulations (Fig. 7). For a given patch-size distribution, the inoculum density determines the range of patch sizes that become populated, and defines the patch size below which most patches are likely to remain unpopulated. This threshold is termed the ‘critical patch size’, marking the tipping point at which reproductive success and *T_50_* begin to decline (∼ 10^5^ µm³ in chip C6HD described in the Results). As demonstrated in Figure 7, our simulations indicate that the inoculum cell density can significantly affect the metapopulation yield. Specifically, varying levels of patchiness and patch-size heterogeneity, result in different amplitudes of metapopulation yield and reproductive success (Fig. 7). These insights have significant implications for microbial ecology and biotechnological applications aiming to either maximize or modulate the yield.

Our experimental results from numerous chips, alongside our computational simulations, indicate that the observed patterns of growth rate, reproductive success, and time to reach the carrying capacity are fairly robust. These patterns appear to be largely independent of the precise shape of growth curves or the specific population growth model applied. The key factor is the density dependent nature of growth rates. Thus, variation in the initial cell densities within patches will significantly influence growth rates and dynamics. While cell seeding governed by processes other than the Poisson distribution can impact initial densities, the key point is that as long as seeding remains stochastic and size-dependent, the resulting initial cell density will significantly affect growth dynamics, which can be anticipated based on the seeding process’s nature. Due to the sub-linear increase in carrying capacity as a function of patch size, reproductive success and the time to reach the carrying capacity tend to peak in intermediate-size patches. The observed deviation from a perfectly linear relationship between droplet volume and the carrying capacity suggests that in our system, factors beyond simple nutrient availability are in play. Possibly, oxygen accessibility, spatial constraints (e.g., on the surface), and potentially other factors, play a role in modulating bacterial growth dynamics in larger droplets, despite having equal nutrient concentration in the suspension sprayed across all droplets. This points at the complex interplay between patch size distribution, carrying capacity, and environmental and biological factors in shaping growth and yield.

We have shown that the division of the metapopulation into growing versus non-growing cells, and its evolution over time, is modulated by the distribution of patch sizes as shown in Figure 4G. This distinction and its dynamics can have important ecological consequences. For example, because growing cells are more susceptible to various antibiotics, survival rates may differ based on the cells’ growth states, which are in turn influenced by patch size. The timing of stress application along the growth dynamics can result in different survival outcomes depending on the distribution of patch size. In the early stages, cells in small patches that have reached their carrying capacity may survive. However, in later stages, cells in both small and large patches may enter the stationary phase, while only those found in intermediate-sized patches continue to grow (Fig. 4G). Additionally, the distribution of patch sizes influences the shift in metapopulation subdivision across patch size ranges over time, generally favoring those in intermediate-sized patches (Supp. Fig. S1).

Our *in vitro* chip, while simplified and focused on clonal single-species populations, does not capture the full complexity of natural microbial habitats, where microbes live in multi-species communities. In our setup, patches were completely isolated, precluding migration that might be common in some natural microbial habitats. Despite these simplifications, the principles we have uncovered offer valuable insights and lay the ground for future studies that may incorporate additional layers of complexity. Furthermore, we believe that this work is valuable for macro ecology and landscape ecology. We suggest that ecology-on-a-chip, using bacteria as a model organism, can be a powerful tool to study fundamental questions in ecology and population dynamics.

In summary, our research shed lights on how natural fragmentation of microbial habitats into micro-patches of various sizes affects microbial growth and population dynamics. This patchiness, observed in environments as diverse as soil porous media, leaf surfaces, and the varied landscape of the human skin and gut, significantly affects microbial growth, reproductive success, and yield, thus affecting the size and composition of microbial communities. Our study demonstrates the critical need to incorporate spatial arrangement into the analysis of samples from natural environments, the experimental designs of microbial studies, and the application of microbiome engineering.

## Materials and Methods

### Bacterial Strain and Growth Conditions

*E. coli* MG-1655 cells harboring the pEB1-mGFPmut2 plasmid where used throughout the study (pEB1-mGFPmut2 was a gift from Philippe Cluzel, Addgene plasmid #103980; http://n2t.net/addgene:103980; RRID: Addgene_103980)^74^. The pEB1-mGFPmut2 plasmid constitutively expresses a green fluorescent protein (GFPmut2) and carries a kanamycin resistance gene. Culture was grown in M9 medium (M9 Minimal Salts Base 5x, Formedium, UK) supplemented with 2 mM MgSO_4_, 0.1 mM CaCl_2_, 20 mM glucose, 40 mM tryptone, 0.1% (v/v) trace metals mixture (1000x Trace Metals Mixture, Teknova) and 50 µg/mL kanamycin. Tryptone-glucose concentrations were set to support a substantial reproductive success on one hand, but avoiding too populated droplets that will complicate cell count on the other hand. Starter cultures were grown over night under agitation (220 rpm) at 37°C. 50 µL from the overnight culture were transferred into 3 mL of fresh medium (same as described above except adjustment of glucose to 10 mM and tryptone to 20 mM) and incubated for an additional 3–5 hr (until OD reached a value of ∼0.5). finally, the OD∼0.5 culture was centrifuge for 5 min at 4300 rcf and the resulting pellet was suspended in 2 mL of fresh medium without kanamycin. The optical density was then adjusted to OD_600_=0.01 or 0.03 and Alexa 647 (Alexa Fluor 647, Invitrogen) was added to the samples at a final concentration of 2 µM.

### µ-SPLASH experiments

µ-SPLASH experiments were conducted in two rounds of experiments (April 2023 [I] and June 2023 [II]) which defers in their technical setting. **Experiment I**: glass-bottom 24-well plates (P24-1.5H-N, cellvis, USA) were used as a carrier vessel for eight µ-SPLASH chips. The peripheral wells (row A; D and column 1; 6) were filled with 2 mL of deionized water (di-H_2_0) and the hollow spaces between the wells were filled with 150 µL di-H_2_0 (except the central hollow space that was left empty) in order to maintain a humid environment. The central wells (B2-5; C2-5) were used to accommodate µ-SPLASH chips, one in each well. A small hole was made in the center of the top plastic lid through which a temperature sensor was inserted. 200 µL of the *E. coli* MG-1655 cell sample (with either OD_600_ of 0.01 or 0.03) were transferred to the loading cup of an airbrush painter (IWATA model HP-M, Japan). The airbrush nozzle was pointed toward the center of a glass bottom well plate through a hollow cylinder (custom printed on an in-house 3D printer; see supplementary for more details) that was inserted to the well. The distance between the airbrush nozzle and the glass surface of the well was approximately 4.5 cm. An air compressor connected to the airbrush was set on 1.4 bar and the fluid adjustment valve of the airbrush was tuned to level 4. A short manual press on the airbrush activation lever was applied to deliver a sprayed sample onto the glass surface of each of the used wells. After all the wells were loaded, the cover lid of the multiwall was assigned and sealed by a stretchable sealing tape on the plate periphery (container seal, CSEAL-75R, USA). The 24-well plate was placed in a stage-top environmental control chamber (H301-K-FRAME, Okolab srl, Italy) equipped with an open frame sample holder (MW-OIL, Okolab srl, Italy) and set to maintain 28 °C. **Experiment II:** the following modification were adopted in the second round of experiments: (1) 6 glass bottom well plate (P6-1.5H-N, cellvis, USA) were used (2) the hollow spaces between the wells were filled with 1.5 mL di-H_2_0 except the smaller hollow space in the corners that were filled with 1 mL di-H_2_0 (3) wells A1-2 and B1-2 were used to accommodate µ-SPLASH chips (two chips in each well) (4) the hollow cylinder used to direct the spray jet had a modified design (5) the 6 well plate was mounted on a 48 well plate holder (48MW, Okolab srl, Italy) inside the stage-top environmental control chamber (the chip were sprayed such that the center of each chip was positioned above the center of an open well of the 48 well plate holder). The described modifications were designated to improve the thermal uniformity between, and within, µ-SPLASH chips.

### Microscopy

Microscopic inspection and image acquisition were performed using an Eclipse Ti-E inverted microscope (Nikon) equipped with Plan Apo 20x/0.75 N.A. air objective. A LED light source (SOLA SE II, Lumencor) was used for fluorescence excitation. GFP fluorescence was excited with a 470/40 filter, and emission was collected with a T495lpxr dichroic mirror and a 525/50 filter. Alexa 647 fluorescence was excited with a 620/60 filter, and emission was collected with a T660lpxr dichroic mirror and a 700/75 filter. Filters and dichroic mirror were purchased from Chroma, USA. A motorized encoded scanning stage (Märzhäuser Wetzlar, DE) was used to image multiple positions of the µ-SPLASH chips. In experiment I, the center of the chip was scanned using the ‘large image mode’ by imaging 12 x 12 adjacent fields of view (with a 5% overlap, 7.64 × 7.64 mm per scan). In experiment II, the center of the chip was scanned using the ‘large image mode’ by imaging 10 x 8 adjacent fields of view (with a 5% overlap, 6.36 × 5.09 mm per scan). Focusing – per field of view–was achieved by using the ‘autofocusing’ mode (step 6 µM; range 18 µM). Images were acquired with an SCMOS camera (ZYLA 19 4.2PLUS, Andor, Oxford Instruments, UK). NIS Elements 5.02 software was used for acquisition.

### Image processing and analysis

Tiff images were exported in 16-bit RGB format as two separate channels: GFP fluorescence and ALEXA fluorescence. Image processing was performed using two open-source software tools: FIJI version 1.53v^75^ and Ilastik version 1.3.3post3^76^. Droplets segmentation was based on the Alexa 647 fluorescence channel while GFP fluorescence channel was used for bacterial cells segmentation.

### Image aligning

To correct fields of views (FOVs) drifts on the XY plane between sequential time points, GFP images were aligned using an automated FIJI Macro script that was developed (script available at 10.6084/m9.figshare.25541083). Initially, large images were cropped to a smaller area (4000 x 2500 pixels) to reduce computational load. The cropped images were then enhanced for better contrast using the FIJI ‘enhanced contrast’ function, and then stacked. A FIJI ‘Linear Stack Alignment with SIFT’ function that identifies key common features across the images was applied. The alignment’s transformation matrix, listing X and Y shifts for each image, was cumulatively applied to ensure consistent alignment of droplet projections over time.

### Droplet’s segmentation

For both experiments (I-II), droplet mask was created based on a single chosen time point. The obtained droplet mask was later used for all time points since droplet area did not change much during the experiment. Tiff images from the Alexa 647 channel were used to create distinct regions of interest (ROIs) for each chip (Fig. 8). Time points t=4 h and t=17 h were selected for Experiment I and II, respectively. The selected images were processed by a FIJI automated script, starting with Gaussian Blur and Median filtering to reduce noise and enhance droplet visibility, followed by conversion to 8-bit files and manual threshold for precise segmentation and finally conversion to binary masks. ‘Fill Holes’ and ‘Watershed’ tools were applied to better define the droplet boundaries. The ‘Analyze Particles’ function in FIJI quantified droplets and produced ZIP (for ROIs) and CSV files, providing the measured area per each droplet in the chip.

**Figure 8.**
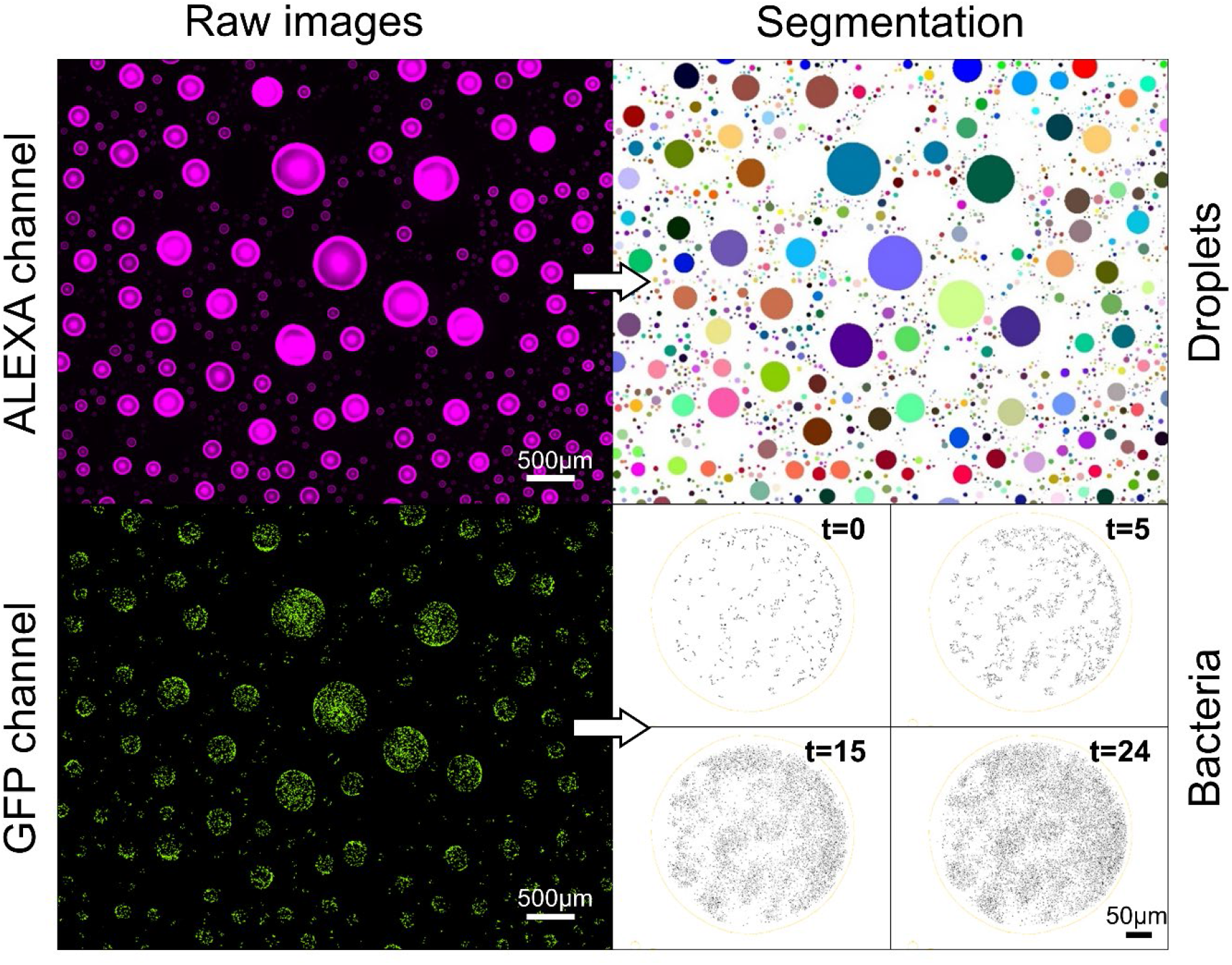
Image processing results. Left: two channels raw data image. Alexa 647 channel captures the droplet boundaries. GFP channel captures bacterial cells. Right: droplets and bacterial segmentations.

### Bacterial segmentation

Single-cell segmentation was achieved by ilastik pixel classification workflow using the GFP channel as a template. Tiff images were converted to HDF5 files using FIJI. Model training was done separately for two time frames to accommodate changes in bacterial organization patterns (single cells vs aggregated cells at later time pointes). Model A was used for the early growth phase (0-7 hours; low density cell), and Model B for the late phase (8-24 hours; high density and aggregates). The models were trained on five selected time points. The training involved manual label of objects classified as cells or background until the model was able to distinguish between all objects and segment individual cells. Post-training, all images were batch processed with the two trained models, and the segmented binary data were exported to Tiff files.

### Counting bacteria within each Droplet ROI

Following droplets and cells segmentation, the droplets ROIs and cells binary data information were merged to assign cell counts per drop per time point (Fig. 9). A FIJI automated script was utilized to generate CSV files summarizing cell counts per droplet in each time point.

### Data pre-processing

Python scripts were employed to merge bacterial count data with the corresponding ROI information for each droplet across the tested chips. For each chip, bacterial counts produced by both Model A and B were recorded. The function integrated ROI data (droplets) with segmented objects (bacterial cells) to accurately assign cell counts to respective droplet. Manual quality check by eye inspection was performed to exclude irregular droplets (less than 5% of all populated droplets).

Droplet size, initially represented by area (µm^2^), was converted to volume. The droplets’ projected area was converted to estimated volume using the following equation (Park *et al.* (2015)^77^):

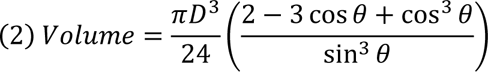

where *D* is the diameter of each droplet and *θ* is the average contact angle of the droplets. Since the contact angle varies between surfaces and depends on the physical properties of the medium, we used a previously measured contact angle of an M9 medium droplet (350 µm in diameter) on a glass surface that was 32° (0.558 rad). Since the contact angle of droplets at our size range (∼50-500 µm in diameter) is not expected to change significantly (according to Park *et al.)*, we could use this previously measured contact angle of 32°.

After converting the area data to volume, weighted counts were calculated for each droplet and time point to extract the most accurate cell count within each droplet according to the two models mentioned before (A and B). Since each model provided its own cell count for each droplet and each time point, but Model A was more accurate for 0-7 h while model B was better for 8-24, the final cell count was calculated from the weighted counts of each model: where model A had higher weight at early times and model B at later times. This was achieved by applying two custom Gaussian functions for the two models: Model A featured µ=0, σ=4; model B featured µ=24, σ=8. The cell counts were the weighted normalization of these weights.

Additional steps are implemented for refining bacterial counts obtained from image processing. The process begins by filtering out droplets that were not populated by cells across all time points. A custom function was then applied to fill in missing data points using the log mean of neighboring values or the nearest valid previous value. The time-series data underwent further processing to adjust for sudden decreases in bacterial counts, applying a log mean fill to smooth these values. LOWESS smoothing is then applied to log-transformed counts to reduce noise.

### Statistical analysis and data modelling

Time-series of single cell data from each droplet was analyzed to calculate population growth dynamics across all 11 chips. For some of the statistical analyses, droplets were binned into groups based on their volume on a logarithmic scale (group ranges were one order of magnitude; 3-4, 4-5, 5-6, 6-7, and 7-8 µm^3^). Statistical differences between the volume bins were examined using permutation tests (1000 permutations each time). Data visualization, modelling and analysis were done using python (Code available at 10.6084/m9.figshare.25541083).

### Computational model and µ-SPLASH simulations

The u-SPLASH simulations were implemented in Phyton. The inputs for the simulation include the total volume of all droplets on the chip, the droplet-size distribution, and the number (or density) of cells at t=0 h. Additionally, a configuration file is utilized to determine the population growth model, the method for determining the carrying capacity in each droplet, and model specific parameters (such as the growth rate ‘*r*’ and other deceleration coefficient ‘*m*’).

In each simulation, cells were randomly assigned to droplets based on their volume, assuming a Poisson process. Growth in each individual droplet is then simulated using the desired model, and growth curve are generated for each droplet. Subsequently, the time-lapse data can be utilized for further downstream data analysis, similar to the approach employed for the experimental data.

Simulation of the C6HD chip presented in the Results, employed a ‘Generalized logistic model’ for growth in each droplet, with parameters derived from the experimental results (specifically, the median values of ‘*r*’ and ‘*m*’). The Carrying capacity (*K*) was determined based on reproductive success, which in turn was inferred from the experimental data using a multiple regression of log initial cell density and log volume, expressed as *RS*=*A* +*B* log_2_ *V* +*C* log_2_*D_0_*. The carrying capacity *K* for each droplet, was then computed based on the calculated RS using the formula *K*=*N_0_*\*2*^RS^*. The exact variables and parameter are listed in Supp. Table. S6.

To explore the effect of patchiness, i.e., the total number of droplets, simulations were conducted with the same total drop volume but partitioned into an increasing number of droplets (ranging from a single drop to 10^4^ droplets. The other parameters were set equals to those fitted to the experimental results of chip C6HD. Further details can be found in Supp. Table. S7. To explore the effect of patch-size heterogeneity chips were simulated with drop size distribution that are log normal with mu that is the mean of drop size across the 11 chips (Supp. Table. S8) and an increasing sigma values.

## Acknowledgements

We thank Jonathan Friedman for valuable comments and discussions and Neta Cohen-Simhi for assistance in imageJ scripts.

## Author contributions

D.M., T.O., and N.K. conceived the study. D.M. and T.O. performed experiments. D.M. performed image processing and data analyses. D.M and N.K. conducted mathematical modeling and simulations. All authors discussed the results and contributed to the final manuscript. N.K. supervised the project. D.M., T.O., and N.K. wrote the manuscript.

## Data and materials availability

Data and code are available at 10.6084/m9.figshare.25541083. All data needed to evaluate the conclusions in the paper are present in the paper and/or the Supplementary Materials.

