## Supplementary materials for "The impact of micro-habitat fragmentation on microbial populations growth dynamics"

**SI Includes:**

**1. Supporting Figures**

**2. Supporting Tables**

### Supplementary Figures

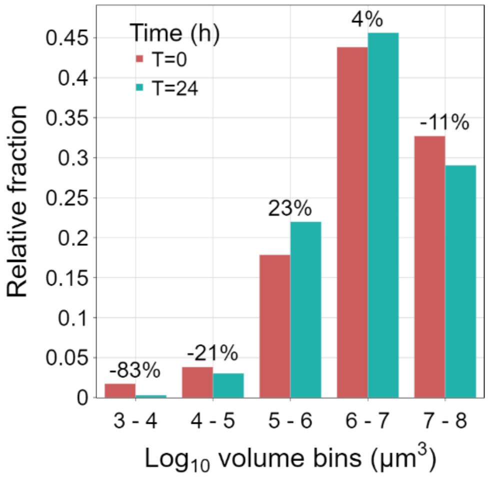


**Figure S1.** The relative change (in %) of bacterial population fraction for different droplet size $\log_{10}$ bins between the initial (t=0 h; red bars) and final (t=24 h; green bars) time points of the experiment. Positive values denote a relative increase in the fraction of the metapopulation residing within that bin, whereas negative values indicate a decrease.

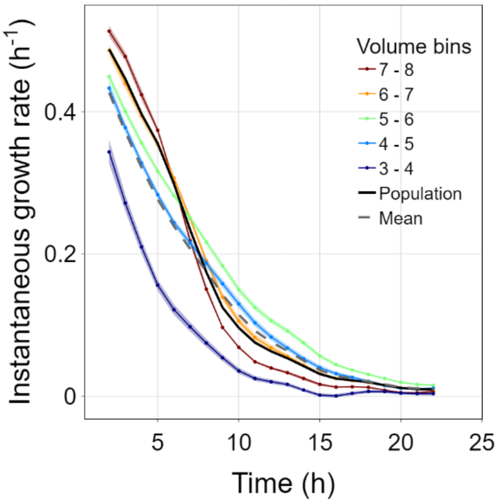


**Figure S2.** The instantaneous growth rate of each volume bin through time of chip C6HD, which extracted from the slope of the curve plotting the natural log of the number of bacterial cells within each droplet over a moving time window of 4 hours.

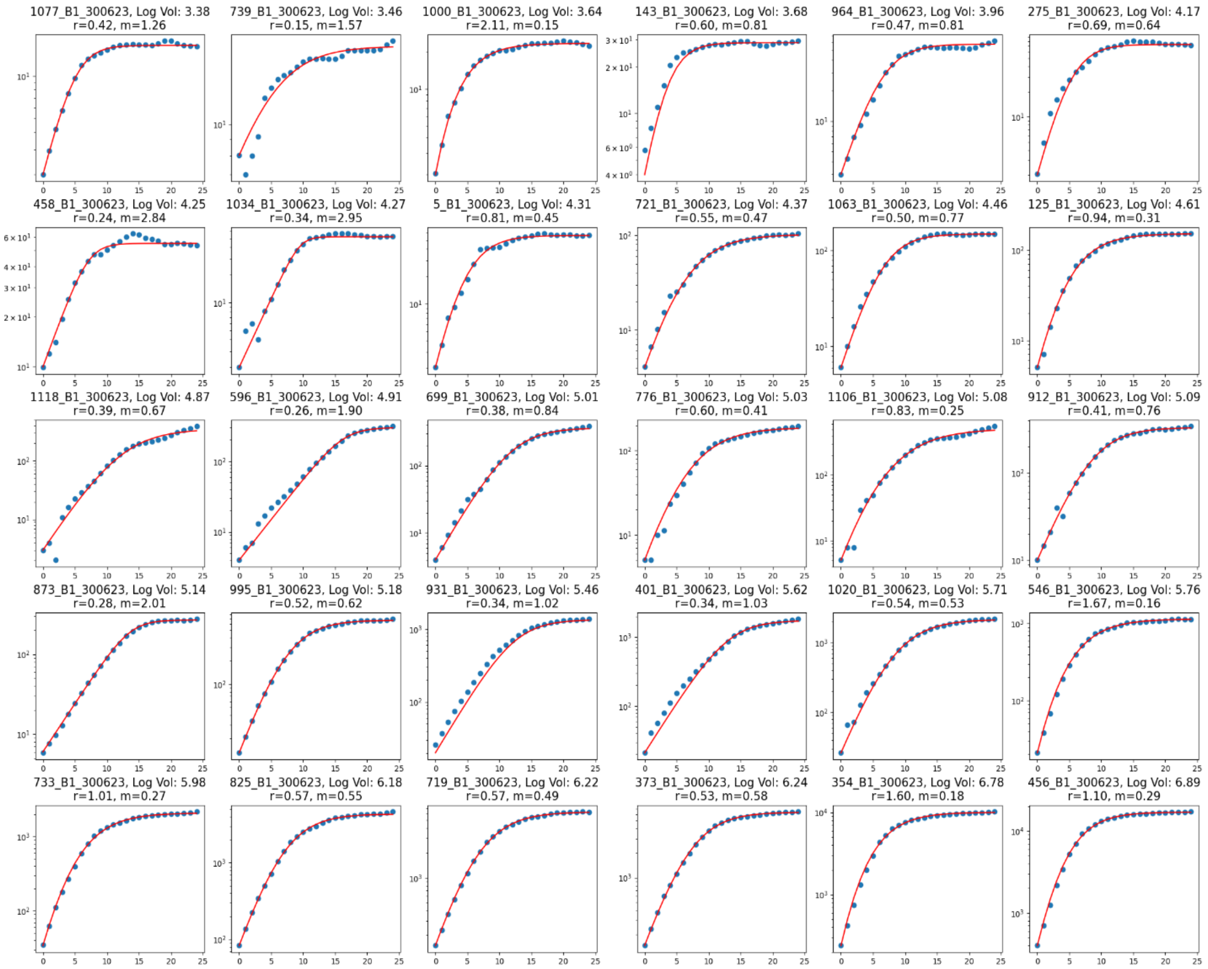


**Figure S3.** Representative sample of growth curves extracted from individual droplets and fitted to a generalized logistic growth model. The blue dots represent the experimental cell count (log-scale) for each time point. The red line represents the fitted model. The sampled subplots are ordered based on the droplets volume from left to right. The fitted model parameters r and m are presented above each plot.

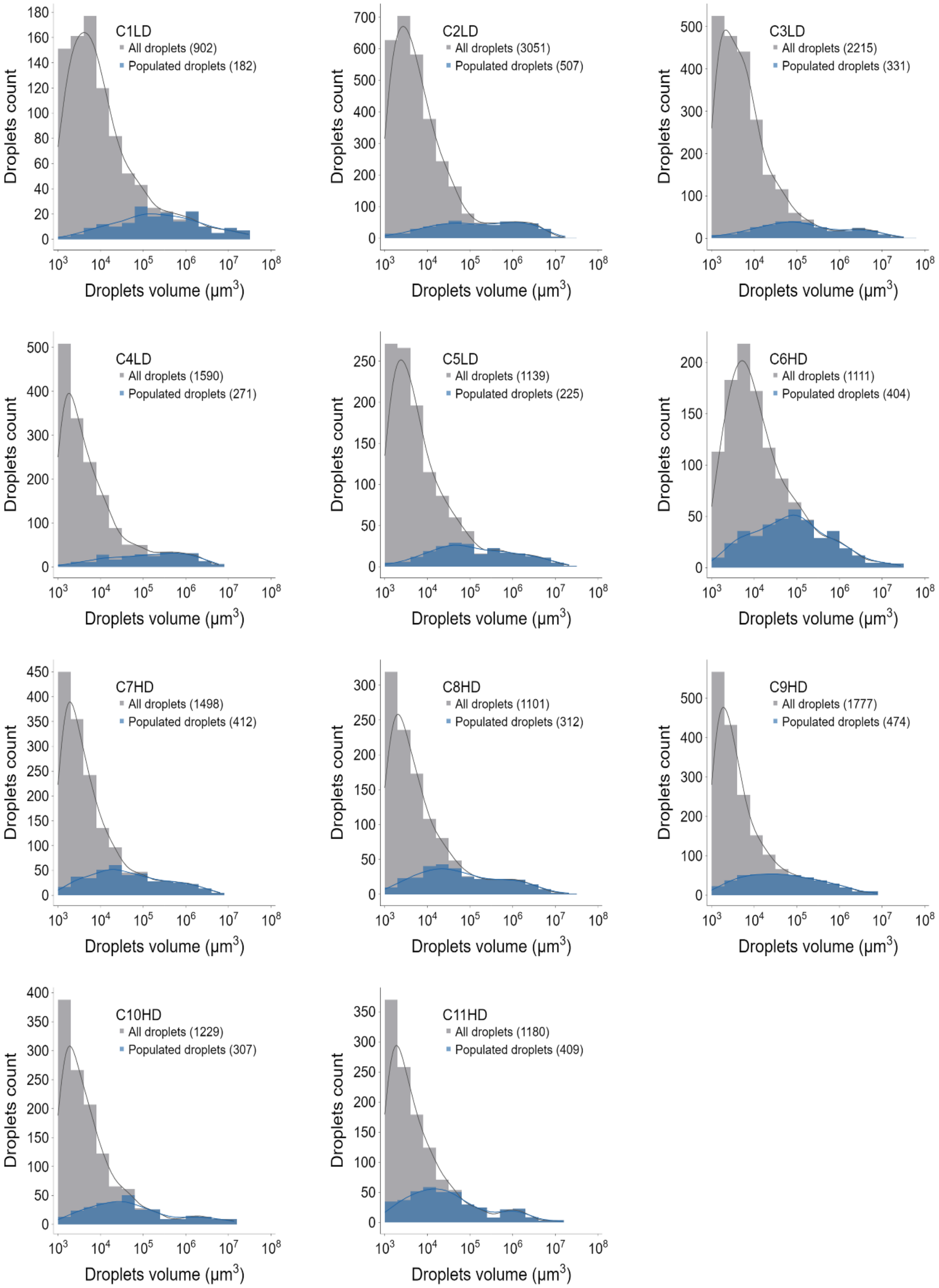


**Figure S4.** Droplets volume distribution across 11 chips (C1-C11; X axis in log scale). Gray bars denote all droplets while blue bars denote bacteria populated droplets.

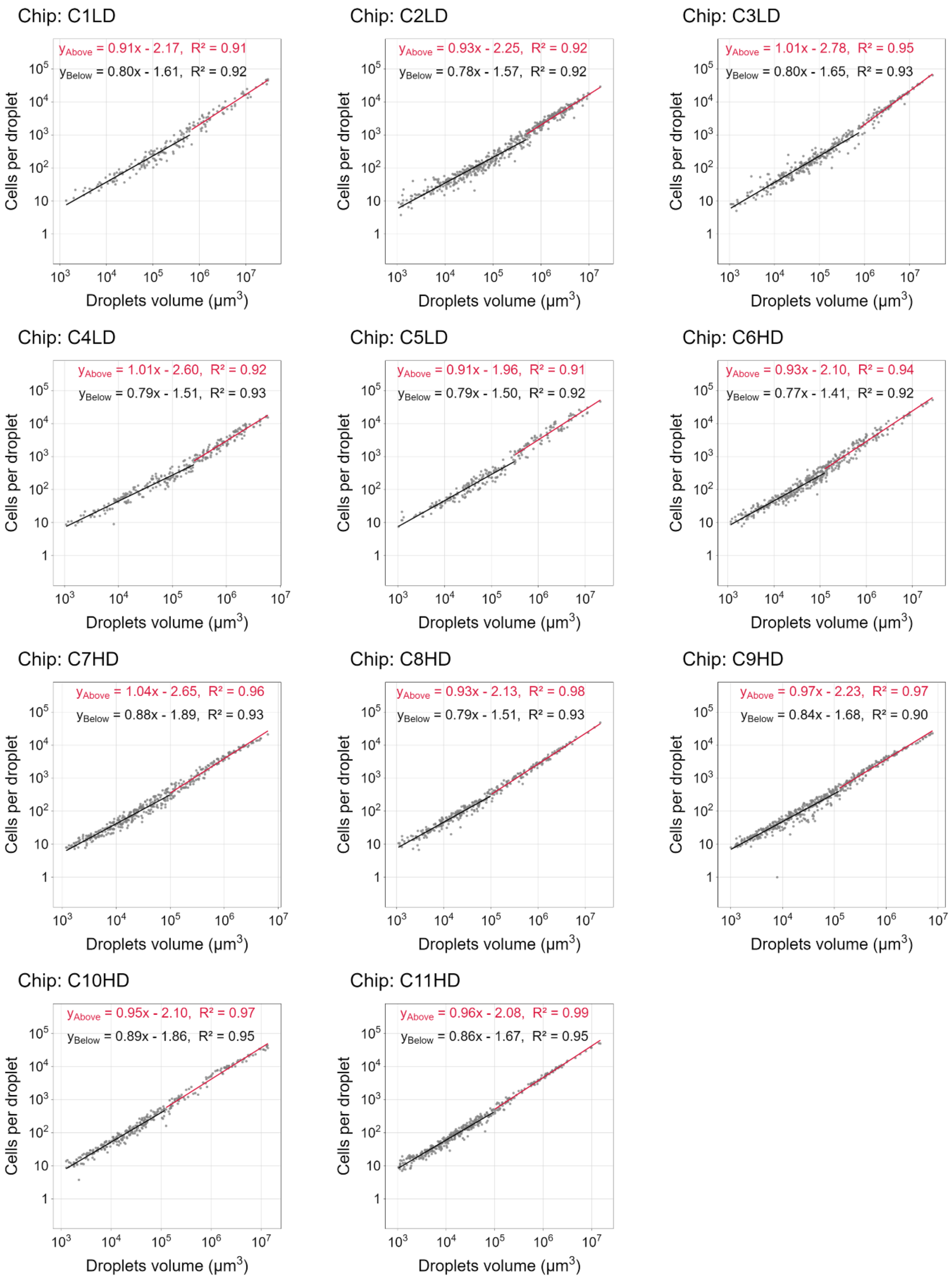


**Figure S5.** The relation between carrying capacity (*K*) and droplet volume (*V*) (log-log plot) across chips. *K* shows two sub-linear phases with different exponents for each chip. The exponents of these two phases are similar across chips.

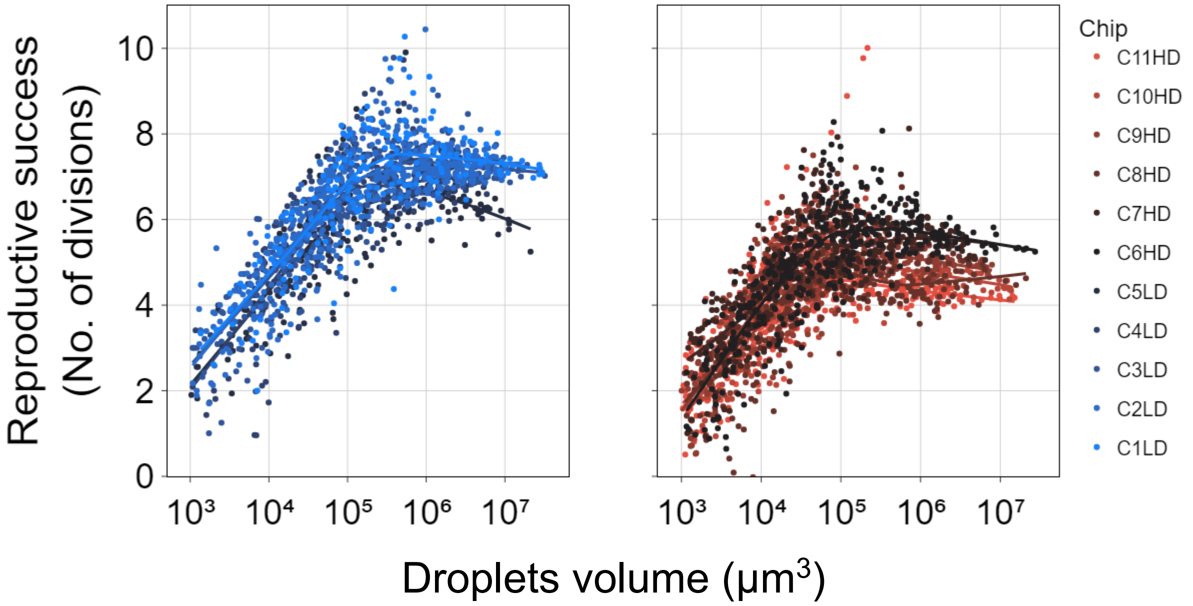


**Figure S6.** Reproductive success (*RS*) across all chips. Left: low density chips (LD; chip's 1-5) in blue. Right: high density chips (HD; chip's 6-11) in red. The colored lines represent the locally weighted scatterplot smoothing (LOWESS, frac=0.4) trendline of each respective chip.

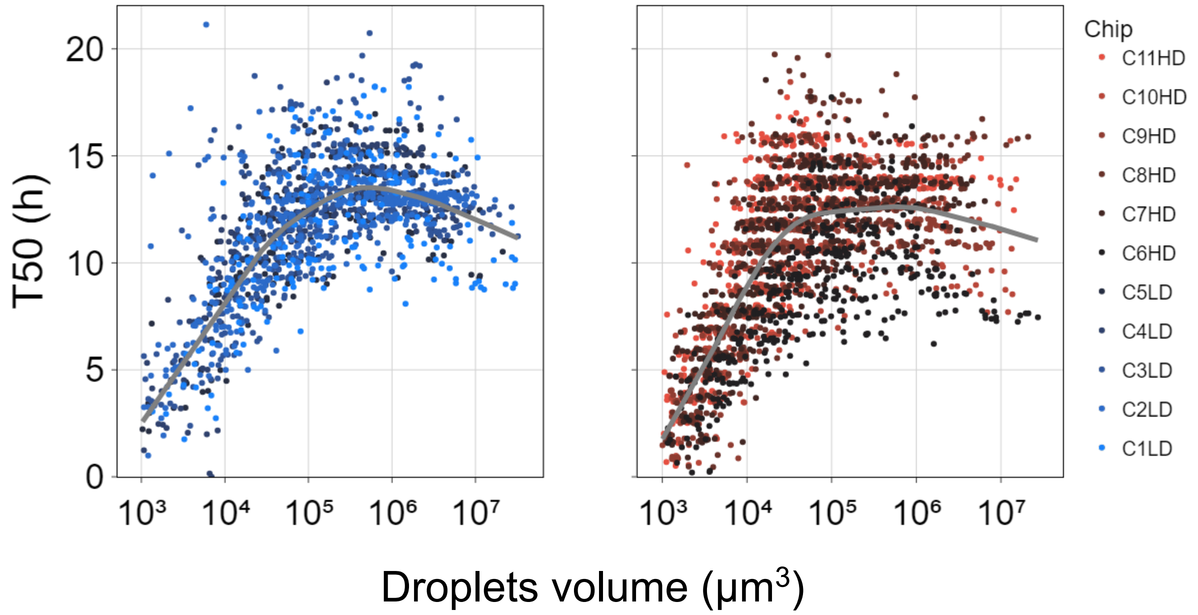


**Figure S7.** Time to reach 50% of carrying capacity ($T_{50}$). Left: low density chips (LD; chip's 1-5) in blue. Right: high density chips (HD; chip's 6-11) in red. The gray line represents the locally weighted scatterplot smoothing (LOWESS, frac=0.4) trendline of all the LD/HD chips respectively.

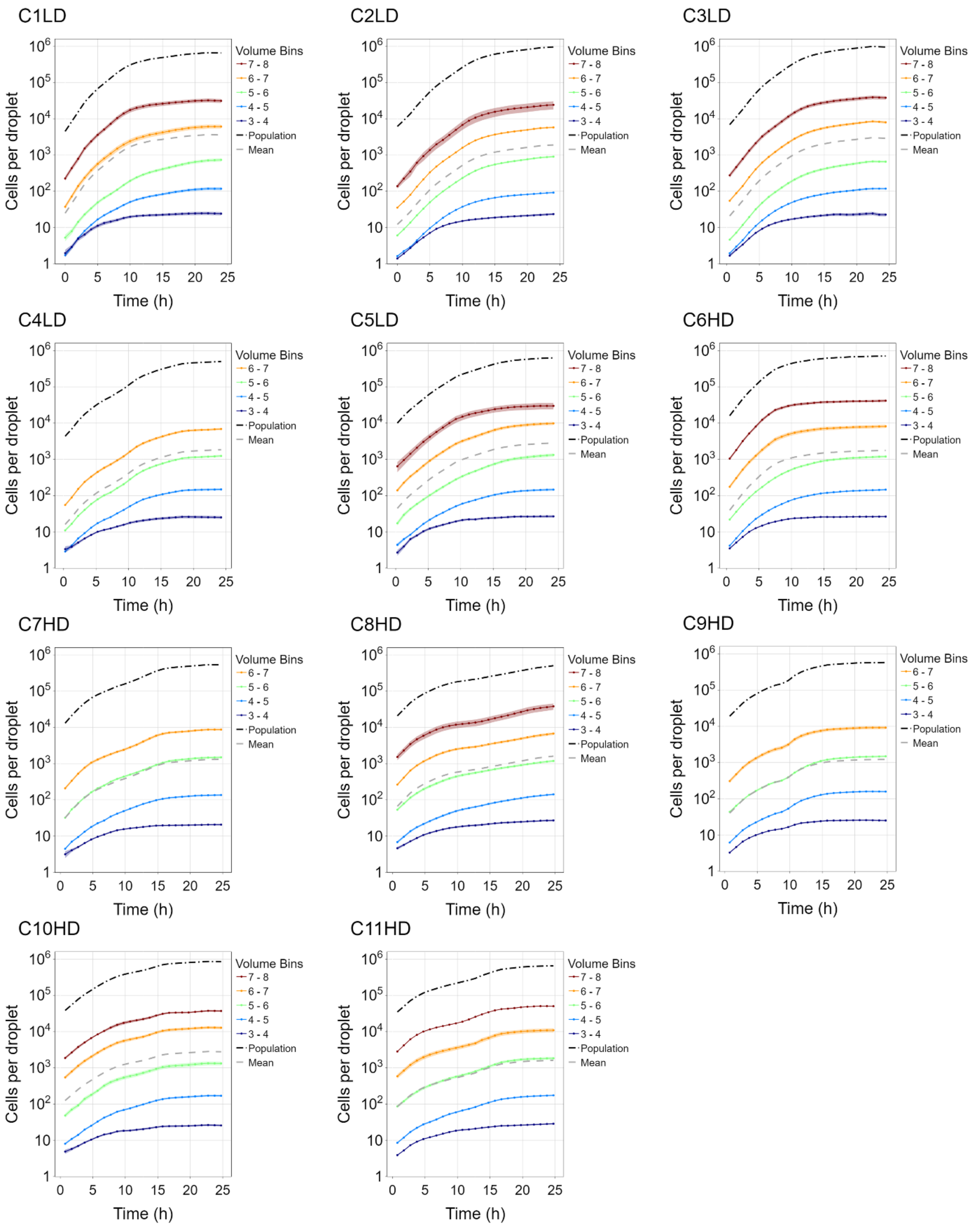


**Figure S8.** Mean growth curves across all chip samples. The colored lines represent the mean cell count per droplet bin volume ($log_{10}$) over time (24 hours). The shaded color of each curve represents the standard error.

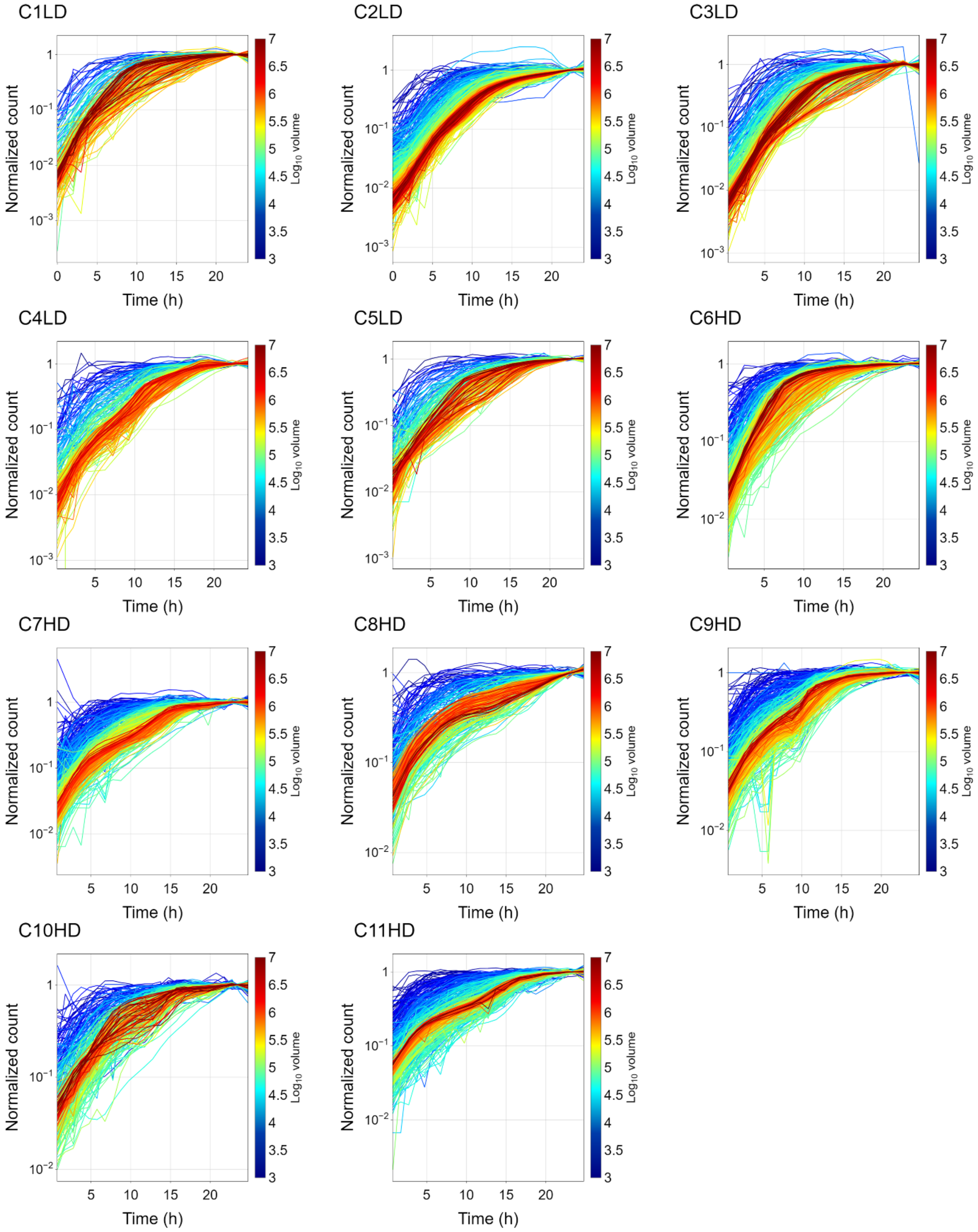


**Figure S9.** Normalized growth curves of each individual droplets across all 11 chips. Normalized growth in each droplet represented by a line colored based on the droplet's volume. Y axis is in log scale.

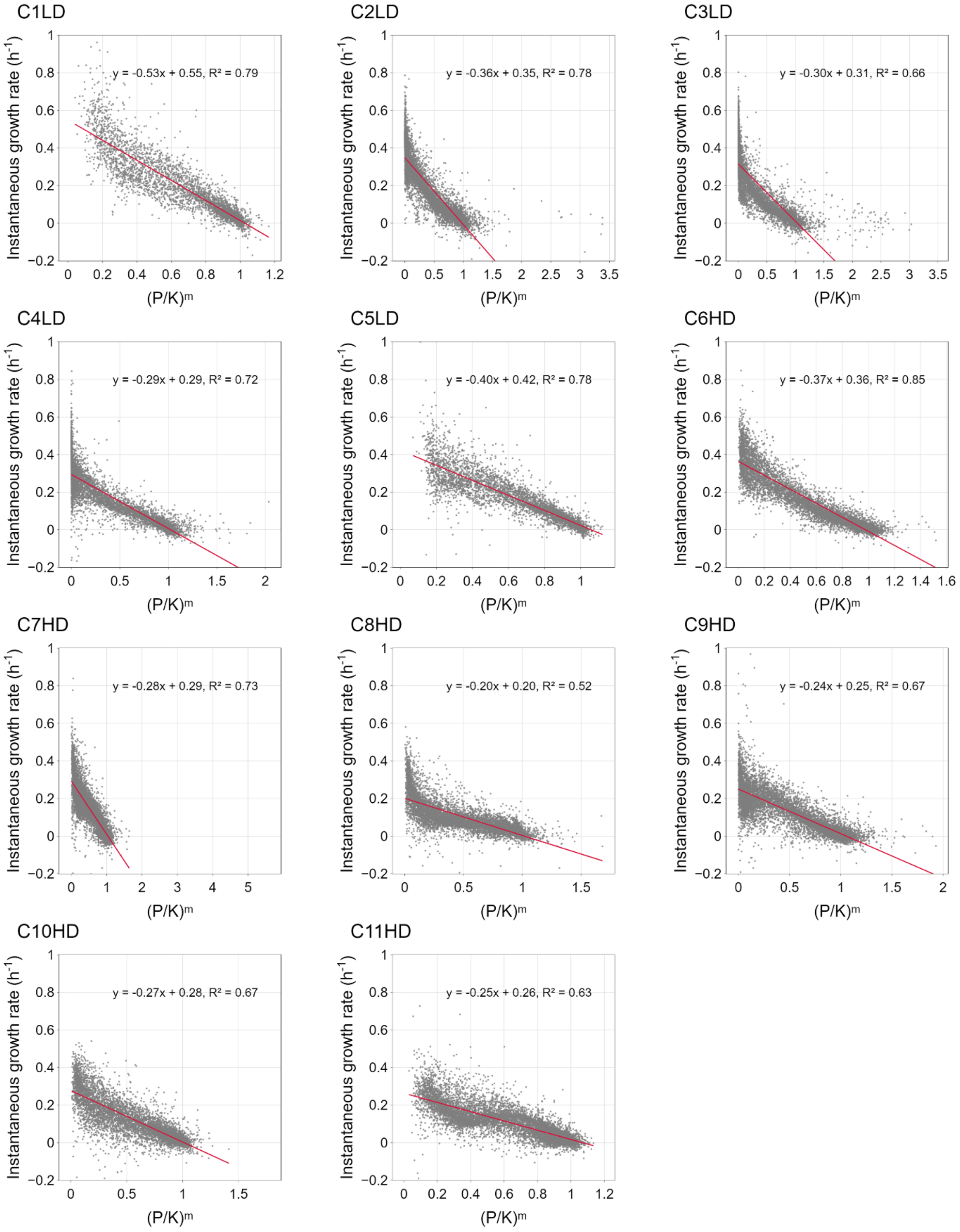


**Figure S10.** Instantaneous growth rates as a function of 'relative density', i.e., the ratio between cell number and carrying capacity in the power of 'm' (m is the fitted deceleration parameter). Mean deceleration parameter of each chips was calculated by averaging the deceleration parameters of all the droplets within that chip. Linear relationship represented by OLS regression line.

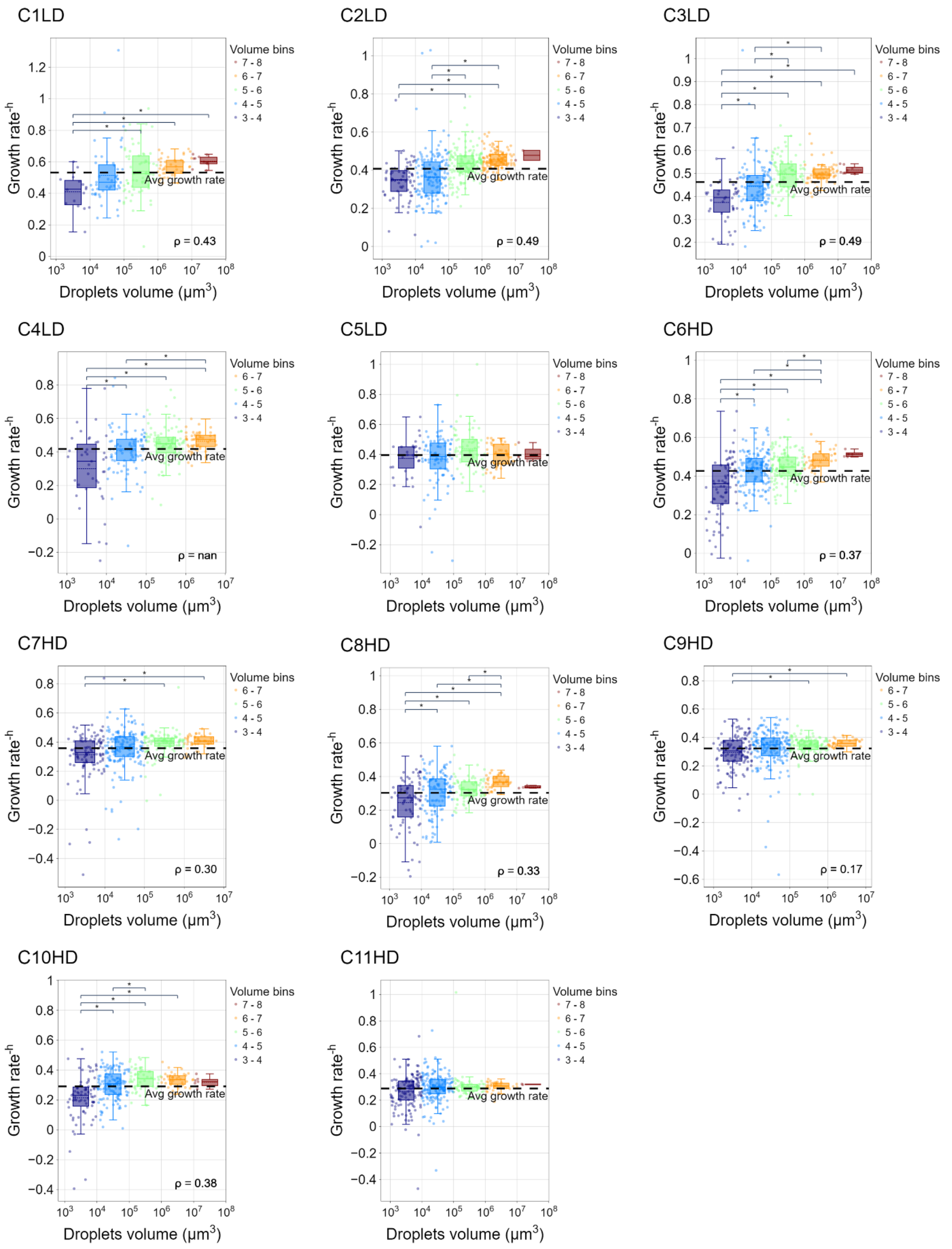


**Figure S11.** Maximal growth rates ($\mu_{max}$) within individual droplets across all 11 chips. The boxplots represent the growth rate distribution of the droplets within each volume bin, colored based on the volume bins. The dashed line represents the overall metapopulation $\mu_{max}$ (of each chip). A permutation tests (n=1000) was used to test statistical significance between the bins' means ($\alpha$=0.01).

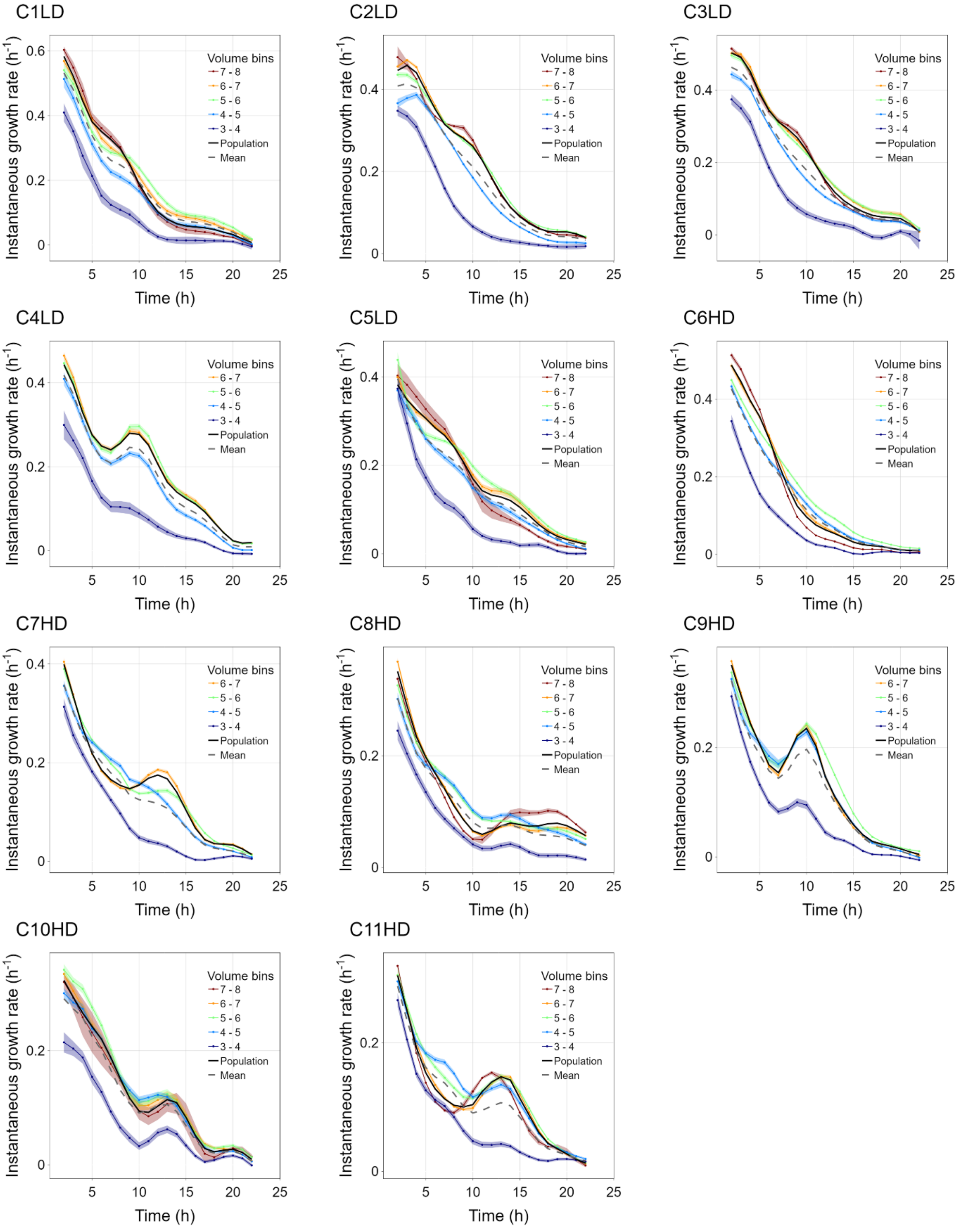


**Figure S12.** The instantaneous growth rate of each volume bin through time and across chips. Rates extracted from the slope of the curve plotting the natural log of the number of bacterial cells within each droplet over a moving time window of 4 hours. The shaded color of each line represents the standard error.

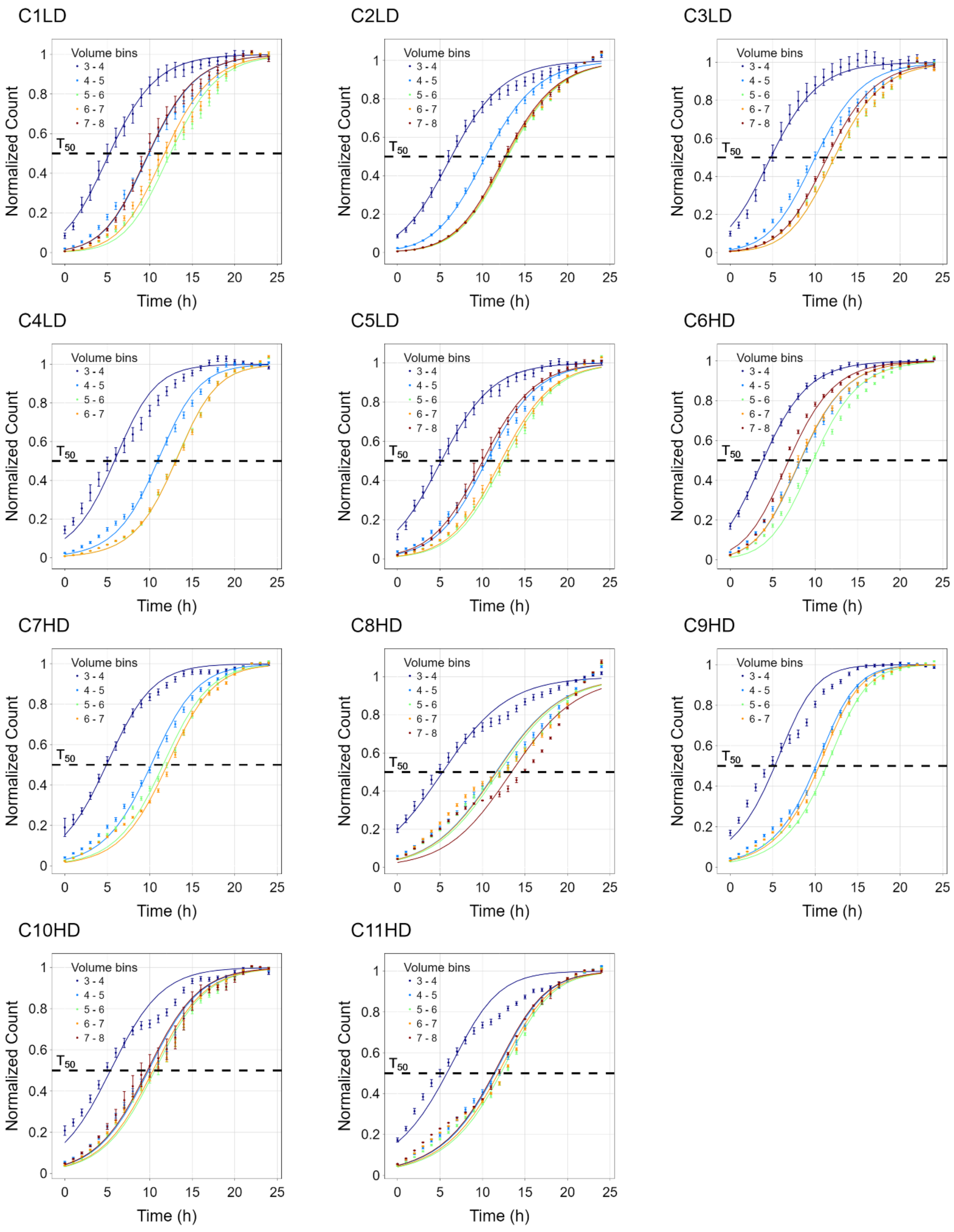


**Figure S13.** Normalized growth curves for each volume bin across chips. Dots and error bars represent Mean ± SE of the experimental data. The line represents the generalized logistic growth model for each volume bin, based on the median r and m values of all droplets, and the mean initial cell density.

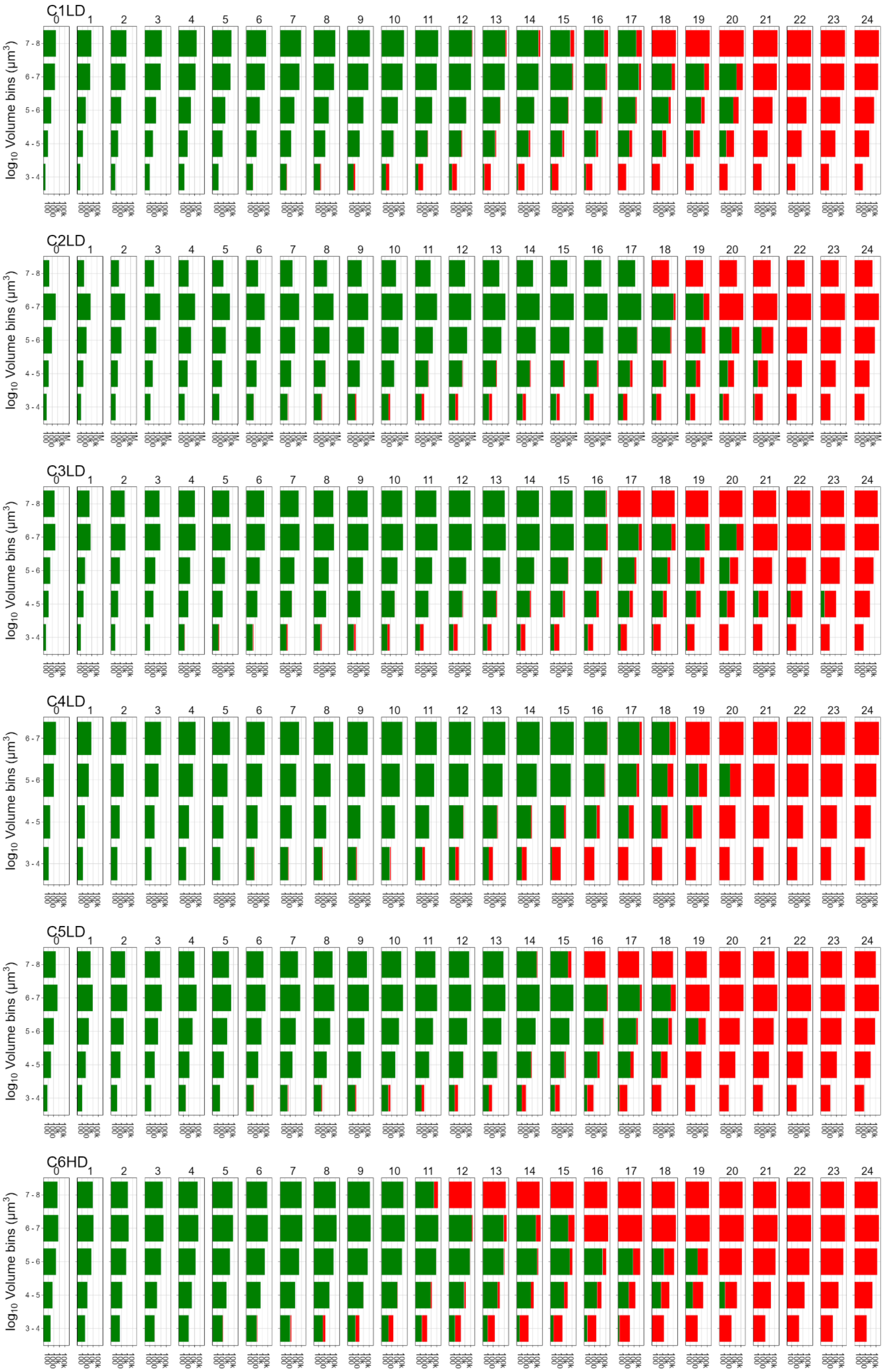
C1LD
C2LD
C3LD
C4LD
C5LD
C6HD
$\log_{10}$ Volume bins ($\mu m^3$)
0 1 2 3 4 5 6 7 8 9 10 11 12 13 14 15 16 17 18 19 20 21 22 23 24
7 - 8
6 - 7
5 - 6
4 - 5
3 - 4

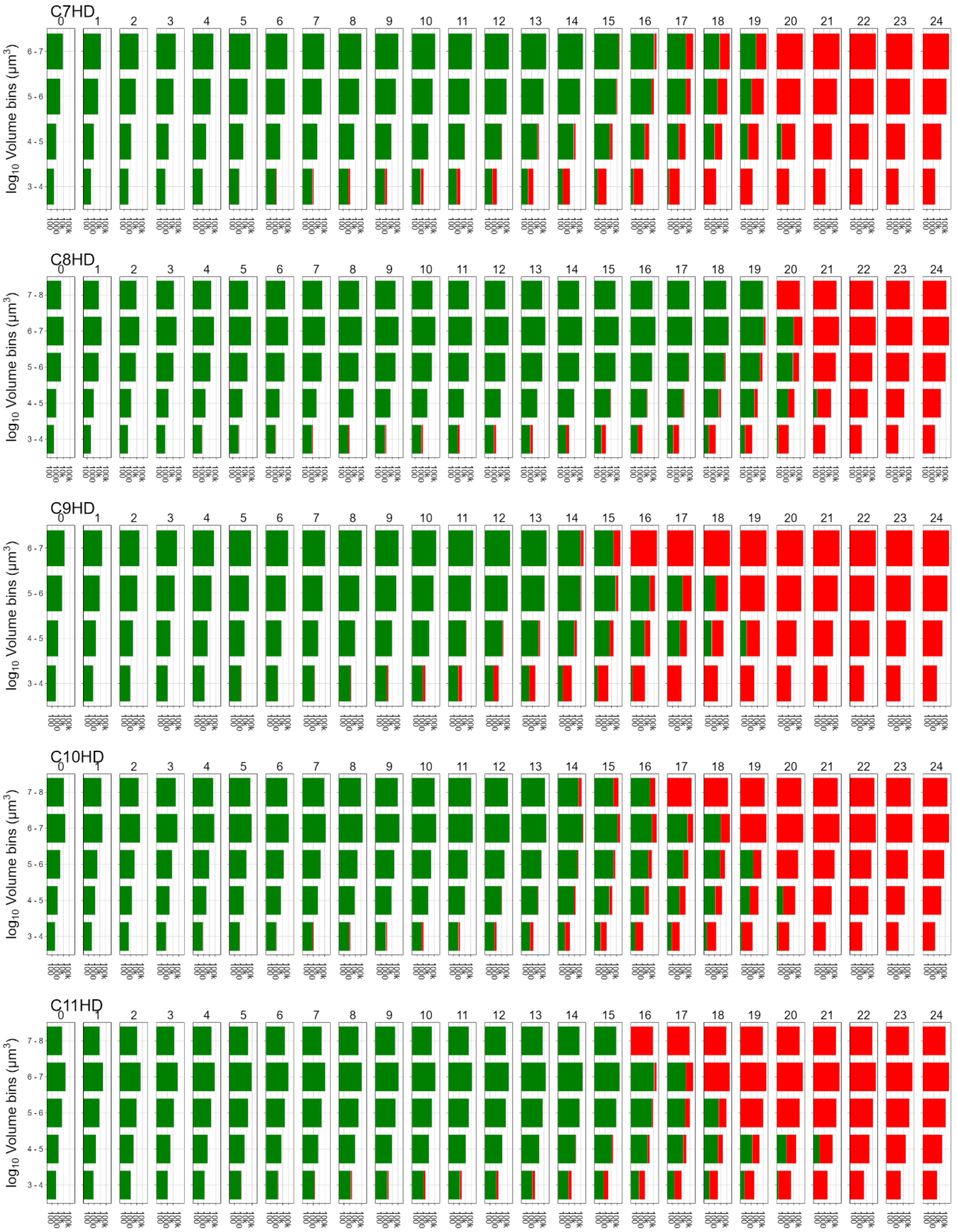


**Figure S14.** Bar chart representing time-lapse of the overall population dynamics and state (growing in GREEN; stationary in RED) over time in each bin, across chips. The bar width represents the cell counts (in log scale) in each drop-volume bin over time.

### Supplementary Tables

| | Coefficient | Std error | t | P>\|t\| |
|---|---|---|---|---|
| **Constant (A)** | -5.0972 | 0.161 | -31.700 | 0.000 |
| **log2_Volume (B)** | -0.0357 | 0.010 | -3.586 | 0.000 |
| **log2_Density (C)** | -0.8108 | 0.018 | -45.387 | 0.000 |

**Table S1**. Coefficients, standard errors, and t-statistic for reproductive success, droplet volume and initial density multiple regression.

| Chip | | All droplet | Populated droplets | Initial cell no. | Final cell number | RS | Overall volume | Initial OD | Initial density | Initial Distribution | |
|---|---|---|---|---|---|---|---|---|---|---|---|
| | | | | | | | | | | mu | sigma |
| A1_30 0623 | C1 LD | 902 | 182 | 4479 | 644844 | 7.17 | 3.94e+08 | 0.01 | 1.1e-5 | 3.75 | 1.231 |
| B3_100 423 | C2 LD | 3051 | 507 | 6475 | 956225 | 7.21 | 5.19e+08 | 0.01 | 1.2e-5 | 3.46 | 1.124 |
| C3_100 423 | C3 LD | 2215 | 331 | 7033 | 955052 | 7.09 | 4.78e+08 | 0.01 | 1.5e-5 | 3.25 | 1.111 |
| A2_26 0623 | C4 LD | 1590 | 271 | 4382 | 505762 | 6.85 | 1.69e+08 | 0.01 | 2.6e-5 | 3.18 | 1.148 |
| A2_25 0623 | C5 LD | 1139 | 225 | 8729 | 629987 | 6.17 | 2.24e+08 | 0.01 | 3.9e-5 | 3.28 | 1.169 |
| B1_300 623 | C6 HD | 1111 | 404 | 15837 | 714708 | 5.5 | 2.88e+08 | 0.03 | 5.5e-5 | 3.69 | 1.153 |
| B2_020 723 | C7 HD | 1498 | 412 | 12408 | 534297 | 5.43 | 1.44e+08 | 0.03 | 8.6e-5 | 3.2 | 1.098 |
| B2_300 623 | C8 HD | 1101 | 312 | 20345 | 498677 | 4.62 | 1.9e+08 | 0.03 | 1.07e-4 | 3.24 | 1.179 |
| B2_260 623 | C9 HD | 1777 | 474 | 18633 | 576445 | 4.95 | 1.69e+08 | 0.03 | 1.1e-4 | 3.18 | 1.084 |
| B2_200 623 | C10 HD | 1229 | 307 | 37365 | 852088 | 4.51 | 2.4e+08 | 0.03 | 1.56e-4 | 3.21 | 1.114 |
| B2_250 623 | C11 HD | 1180 | 409 | 34158 | 656372 | 4.26 | 1.54e+08 | 0.03 | 2.22e-4 | 3.20 | 1.103 |

**Table S2.** Cell numbers and droplet data from all 11 chips

| Chip | Coefficient | | Std error | | t | | P>\|t\| | | $R^2$ | |
|---|---|---|---|---|---|---|---|---|---|---|
| | Initial cell count | Carrying capacity | Initial cell count | Carrying capacity | Initial cell count | Carrying capacity | Initial cell count | Carrying capacity | Initial cell count | Carrying capacity |
| C1LD | 0.89 | 0.88 | 0.04 | 0.01 | 19.98 | 74.18 | 1.87E-33 | 7.06E-137 | 0.83 | 0.97 |
| C2LD | 0.85 | 0.88 | 0.03 | 0.01 | 31.4 | 129.3 | 1.87E-82 | 0 | 0.82 | 0.97 |
| C3LD | 0.96 | 0.9 | 0.03 | 0.01 | 31.89 | 105.1 | 1.04E-57 | 3.35E-255 | 0.9 | 0.97 |
| C4LD | 0.92 | 0.91 | 0.05 | 0.01 | 18.1 | 97.32 | 1.77E-34 | 1.06E-211 | 0.75 | 0.97 |
| C5LD | 1 | 0.91 | 0.04 | 0.01 | 22.27 | 88.24 | 1.53E-35 | 2.10E-175 | 0.86 | 0.97 |
| C6HD | 0.96 | 0.88 | 0.03 | 0.01 | 30.17 | 115.8 | 8.85E-51 | 8.15e-311 | 0.9 | 0.97 |
| C7HD | 1.03 | 0.96 | 0.03 | 0.01 | 30.17 | 145.4 | 7.47E-48 | 0 | 0.91 | 0.98 |
| C8HD | 0.9 | 0.87 | 0.04 | 0.01 | 20.73 | 140.3 | 1.07E-33 | 1.68E-282 | 0.84 | 0.98 |
| C9HD | 0.97 | 0.92 | 0.02 | 0.01 | 39.56 | 127.5 | 5.71E-60 | 0 | 0.94 | 0.97 |
| C10HD | 1 | 0.94 | 0.03 | 0.01 | 31.95 | 145.5 | 6.08E-39 | 7.28E-284 | 0.95 | 0.99 |
| C11HD | 1.04 | 0.92 | 0.02 | 0.01 | 54.8 | 177.2 | 1.68E-55 | 0 | 0.98 | 0.99 |

**Table S3.** Summary of Ordinary Least Squares Regression analysis for initial and final cell counts across 11chips.

| Chip | Vc | **Coefficient** | | **Std error** | | **t** | | **P>\|t\|** | | **R2** | |
|---|---|---|---|---|---|---|---|---|---|---|---|
| | | Phase 1 | Phase 2 | Phase 1 | Phase 2 | Phase 1 | Phase 2 | Phase 1 | Phase 2 | Phase 1 | Phase 2 |
| C1LD | 5.8 | 0.80 | 0.91 | 0.02 | 0.04 | 38.15 | 24.24 | 1.16E-69 | 2.25E-31 | 0.92 | 0.91 |
| C2LD | 5.7 | 0.78 | 0.93 | 0.01 | 0.02 | 60.34 | 44.55 | 1.52E-176 | 1.65E-100 | 0.92 | 0.92 |
| C3LD | 5.9 | 0.80 | 1.01 | 0.01 | 0.03 | 55.40 | 37.47 | 6.18E-141 | 6.30E-53 | 0.93 | 0.95 |
| C4LD | 5.4 | 0.79 | 1.01 | 0.02 | 0.03 | 44.37 | 36.63 | 1.63E-88 | 1.36E-65 | 0.93 | 0.92 |
| C5LD | 5.5 | 0.79 | 0.91 | 0.02 | 0.03 | 40.36 | 28.73 | 4.85E-80 | 3.08E-43 | 0.92 | 0.91 |
| C6HD | 5.1 | 0.77 | 0.93 | 0.01 | 0.02 | 52.77 | 48.19 | 3.75E-136 | 8.66E-95 | 0.92 | 0.94 |
| C7HD | 5 | 0.88 | 1.04 | 0.01 | 0.02 | 59.31 | 61.40 | 1.28E-154 | 1.26E-103 | 0.93 | 0.96 |
| C8HD | 5 | 0.79 | 0.93 | 0.02 | 0.01 | 49.20 | 73.21 | 1.99E-110 | 1.44E-99 | 0.93 | 0.98 |
| C9HD | 5.1 | 0.84 | 0.97 | 0.02 | 0.01 | 53.22 | 66.47 | 2.56E-161 | 1.19E-112 | 0.90 | 0.97 |
| C10HD | 5.1 | 0.89 | 0.95 | 0.01 | 0.02 | 62.43 | 50.97 | 2.53E-139 | 1.21E-67 | 0.95 | 0.97 |
| C11HD | 5 | 0.86 | 0.96 | 0.01 | 0.01 | 77.97 | 85.98 | 4.97E-203 | 2.82E-96 | 0.95 | 0.99 |

**Table S4.** Carrying capacity **(***K***)** as a function of droplet volume (*V*) show two phases of sub-linear relation.

| Chip | Parameter | Coefficient | Std error | t | P>\|t\| |
|---|---|---|---|---|---|
| C1LD | Constant (A) | -4.778579 | 0.270509 | -17.665155 | 4.43E-41 |
| | $\log_2$_Volume (B) | -0.021394 | 0.01427 | -1.499262 | 1.36E-01 |
| | $\log_2$_Density (C) | -0.76594 | 0.025249 | -30.334885 | 1.85E-72 |
| C2LD | Constant (A) | -4.680549 | 0.153675 | -30.457494 | 2.43E-116 |
| | $\log_2$_Volume (B) | 0.010667 | 0.009459 | 1.127668 | 2.60E-01 |
| | $\log_2$_Density (C) | -0.718123 | 0.016798 | -42.749833 | 9.32E-170 |
| C3LD | Constant (A) | -4.855728 | 0.186442 | -26.044124 | 7.67E-82 |
| | $\log_2$_Volume (B) | 0.016657 | 0.010596 | 1.571993 | 1.17E-01 |
| | $\log_2$_Density (C) | -0.727482 | 0.019141 | -38.005565 | 3.36E-122 |
| C4LD | Constant (A) | -5.484374 | 0.186317 | -29.435783 | 5.94E-86 |
| | $\log_2$_Volume (B) | 0.009019 | 0.012812 | 0.703997 | 4.82E-01 |
| | $\log_2$_Density (C) | -0.795902 | 0.0207 | -38.450074 | 4.42E-111 |
| C5LD | Constant (A) | -5.038812 | 0.236391 | -21.315591 | 1.29E-55 |
| | $\log_2$_Volume (B) | -0.00377 | 0.012328 | -0.305815 | 7.60E-01 |
| | $\log_2$_Density (C) | -0.778164 | 0.023674 | -32.870566 | 2.98E-87 |
| C6HD | Constant (A) | -5.097174 | 0.160795 | -31.699891 | 2.75E-111 |
| | $\log_2$_Volume (B) | -0.035683 | 0.009951 | -3.585792 | 3.78E-04 |
| | $\log_2$_Density (C) | -0.81081 | 0.017864 | -45.38714 | 4.48E-160 |
| C7HD | Constant (A) | -5.944518 | 0.170932 | -34.777005 | 3.13E-124 |
| | $\log_2$_Volume (B) | 0.020344 | 0.007861 | 2.58797 | 1.00E-02 |
| | $\log_2$_Density (C) | -0.816115 | 0.017564 | -46.465744 | 2.93E-165 |
| C8HD | Constant (A) | -4.856769 | 0.139581 | -34.79534 | 6.64E-109 |
| | $\log_2$_Volume (B) | -0.071935 | 0.00721 | -9.97688 | 1.67E-20 |

| | | | | | |
|---|---|---|---|---|---|
| | $log_2$_Density (C) | -0.828011 | 0.015467 | -53.534418 | 2.29E-158 |
| C9HD | Constant (A) | -5.162353 | 0.162532 | -31.76205 | 3.63E-119 |
| | $log_2$_Volume (B) | -0.003007 | 0.008982 | -0.334716 | 7.38E-01 |
| | $log_2$_Density (C) | -0.784406 | 0.01819 | -43.122984 | 1.18E-165 |
| C10HD | Constant (A) | -5.59527 | 0.167821 | -33.34067 | 1.47E-103 |
| | $log_2$_Volume (B) | -0.013102 | 0.007365 | -1.779047 | 7.62E-02 |
| | $log_2$_Density (C) | -0.837393 | 0.01784 | -46.93954 | 2.54E-141 |
| C11HD | Constant (A) | -5.33235 | 0.11138 | -47.875095 | 4.58E-169 |
| | $log_2$_Volume (B) | -0.035112 | 0.005512 | -6.370473 | 5.11E-10 |
| | $log_2$_Density (C) | -0.858319 | 0.011852 | -72.417662 | 2.98E-234 |

**Table S5.** Multiple Linear Regression coefficients across chips.

| Variable | Value | Comments |
|---|---|---|
| Total drop volume | $2.88*10^8$ µ$m^3$ | As in chip C6HD |
| Number of cells at t=0 | 15850 cells | Per chip |
| Droplet distribution | Log normal | Similar to chip C6HD<br>Mean=$10^{4.15}$, Sigma =$10^{1.2}$ |
| Reproductive success relation | RS=a+b $log_2$(drop_sz) + c $log_2$(drop_$N_0$/drop_sz) | As in chip C6HD experimental results<br>a=-5.1<br>b= - 0.0357<br>c=0.81; |
| Simulation time frame | 0-24 hours | |

**Table. S6** Simulation settings and parameters in the simulation of chip C6HD as presented in the Results.

| Variable | Value | Comments |
|---|---|---|
| Total drop volume | $10^8$ µm$^3$ | As in chip C6HD |
| Number of cells at t=0 | 100,1000,10000, 100000 | For each set of drop-size simulation tested three initial cell numbers. |
| Droplet distribution i (Patchiness) | Droplets were of equal volume | Droplet size range (in µm$^3$): $10^4$,$10^{4.5}$,$10^5$,$10^{5.5}$,$10^6$,$10^{6.5}$,$10^7$,$10^{7.5}$,$10^8$. |
| Droplet distribution ii (Patch-size heterogeneity) | Droplets followed log-normal distribution with mu=10^3.33 µm$^3$ | Sigma range: 1.1, 1.2, 1.5, 2, 3, 5, 10, 20, 50, 100 |
| Reproductive success relation | RS= a + b*$\log_2$(drop_sz) + c*$\log_2$(drop_N0/drop_sz) | As in chip C6HD experimental results a=-5.1 b= - 0.0357 c=0.81; |
| Simulation time frame | 0-24 hours | |

**Table S7.** Simulation settings and parameters exploring the impact of patchiness and patch-size heterogeneity on population dynamics.

| Chip | mu | sigma |
|---|---|---|
| C1LD | 3.75 | 1.064 |
| C2LD | 3.46 | 1.015 |
| C3LD | 3.25 | 1.045 |
| C4LD | 3.18 | 1.084 |
| C5LD | 3.28 | 1.095 |
| C6HD | 3.69 | 1.044 |
| C7HD | 3.2 | 1.035 |
| C8HD | 3.24 | 1.106 |
| C9HD | 3.18 | 1.025 |
| C10HD | 3.21 | 1.044 |
| C11HD | 3.20 | 1.037 |

**Table S8.** The estimated mu and sigma of the log-transformed droplet volume distribution for each chip.
